# An Atlas of Human and Murine Genetic Influences on Osteoporosis

**DOI:** 10.1101/338863

**Authors:** John A. Morris, John P. Kemp, Scott E. Youlten, Laetitia Laurent, John G. Logan, Ryan Chai, Nicholas A. Vulpescu, Vincenzo Forgetta, Aaron Kleinman, Sindhu Mohanty, C. Marcelo Sergio, Julian Quinn, Loan Nguyen-Yamamoto, Aimee Lee Luco, Jinchu Vijay, Marie-Michelle Simon, Albena Pramatarova, Carolina Medina-Gomez, Katerina Trajanoska, Elena J. Ghirardello, Natalie C. Butterfield, Katharine F. Curry, Victoria D. Leitch, Penny C. Sparkes, Anne-Tounsia Adoum, Naila S. Mannan, Davide Komla-Ebri, Andrea S. Pollard, Hannah F. Dewhurst, Thomas Hassall, Michael-John G Beltejar, Douglas J Adams, Suzanne M. Vaillancourt, Stephen Kaptoge, Paul Baldock, Cyrus Cooper, Jonathan Reeve, Evangelia Ntzani, Evangelos Evangelou, Claes Ohlsson, David Karasik, Fernando Rivadeneira, Douglas P. Kiel, Jonathan H. Tobias, Celia L. Gregson, Nicholas C. Harvey, Elin Grundberg, David Goltzman, David J. Adams, Christopher J. Lelliott, David A. Hinds, Cheryl L. Ackert-Bicknell, Yi-Hsiang Hsu, Matthew T. Maurano, Peter I. Croucher, Graham R. Williams, J. H. Duncan Bassett, David M. Evans, J. Brent Richards

## Abstract

Osteoporosis is a common debilitating chronic disease diagnosed primarily using bone mineral density (BMD). We undertook a comprehensive assessment of human genetic determinants of bone density in 426,824 individuals, identifying a total of 518 genome-wide significant loci, (301 novel), explaining 20% of the total variance in BMD—as estimated by heel quantitative ultrasound (eBMD). Next, meta-analysis identified 13 bone fracture loci in ~1.2M individuals, which were also associated with BMD. We then identified target genes from cell-specific genomic landscape features, including chromatin conformation and accessible chromatin sites, that were strongly enriched for genes known to influence bone density and strength (maximum odds ratio = 58, P = 10^−75^). We next performed rapid throughput skeletal phenotyping of 126 knockout mice lacking eBMD Target Genes and showed that these mice had an increased frequency of abnormal skeletal phenotypes compared to 526 unselected lines (P < 0.0001). In-depth analysis of one such Target Gene, *DAAM2*, showed a disproportionate decrease in bone strength relative to mineralization. This comprehensive human and murine genetic atlas provides empirical evidence testing how to link associated SNPs to causal genes, offers new insights into osteoporosis pathophysiology and highlights opportunities for drug development.

## Introduction

Osteoporosis is a common, aging-related disease characterized by decreased bone strength and consequent increased risk of fracture.^1^ Bone mineral density (BMD), the most clinically relevant risk factor when diagnosing osteoporosis, is highly heritable^2^ and is a strong risk factor for fracture.^3^ While there have been no large-scale genome-wide association studies (GWAS) for fracture to date, previous GWAS for BMD have demonstrated that BMD is a highly polygenic trait.^2^ Recently, we identified 203 loci associated with estimated BMD by measuring quantitative heel ultrasound (eBMD), explaining 12% of its variance, demonstrating this polygenicity.^4^

eBMD is predictive of fracture and is highly heritable (50-80%).^5–9^ While BMD measured from dual-energy X-ray absorptiometry (DXA)-scanning is most often used in clinical settings, our recent GWAS for eBMD identified 84% of all currently known genome-wide significant loci for DXA-BMD^4^ and effect sizes were concordant between the two traits (Pearson’s *r* = 0.69 for lumbar spine and 0.64 for femoral neck).^4^ The largest GWAS to date for DXA-derived BMD measures contained only 66,628 individuals.^10^ Both ultrasound and DXA-derived BMD are strongly associated with fracture risk where a standard deviation decrease in either metric is associated with approximately a ~1.5-fold increase in the risk of osteoporotic fracture,^3,11^ and both traits are highly polygenic.

Little is known about how to reliably map associated genomic loci to their causal genes. However, highly polygenic traits such as bone density offer the opportunity to empirically test which methods link associated SNPs to genes enriched for causal proteins. Causal proteins can be identified in human clinical trials when their manipulation by medications leads to changes in BMD.^2^ Another source of causal proteins is Mendelian genetic conditions, which may constitute human knockouts and can also strongly implicate key genes that underlie bone physiology.^12^ Given a sufficient number of associated loci, the different genomic characteristics that link a SNP to these causal proteins can be tested. These include genomic landscape characteristics such as cell-specific 3-dimensional (3D) contact domains, cell-specific open chromatin states, physical proximity and the presence of coding variation. Furthermore, samples from knockout mice generated by large-scale programs, such as the International Knockout Mouse Consortium (IKMC), can be used to identify genes whose deletion results in an abnormal skeletal phenotype. This rapid-throughput phenotyping data can then be used to determine whether outlier bone phenotypes are enriched in mice harboring deletions of genes identified by GWAS in humans.

Here, we present the most comprehensive investigation of human and murine genetic influences on bone density and fracture to date. We not only undertook a GWAS of 426,824 individuals for eBMD in the UK Biobank, explaining 20% of its variance and identifying 301 novel loci, but also identified the genetic determinants of fracture in up to 1.2 million individuals combining the UK Biobank and 23andMe cohorts. We then assessed the SNP-level and genomic landscape characteristics that mapped associated SNPs to genes that were enriched for known bone density proteins. We identified Target Genes that were enriched up to 58-fold for known causal genes and for genes differentially expressed in *in vivo* osteocytes compared to bone marrow cell models. Finally, we investigated whether deletion of GWAS-identified genes resulted in skeletal abnormalities *in vivo* by undertaking rapid-throughput phenotyping of knockout mice, which included 126 Target Genes. Mice harboring deletions of these 126 Target Genes were strongly enriched for outlier skeletal phenotypes. A convergence of human genetic, murine genetic, *in vivo* bone-cell expression and *in vitro* cell culture data all pointed to a role for *DAAM2* in osteoporosis. This was further investigated by detailed analysis of mice with a hypomorphic allele of *Daam2. Daam2* knockdown resulted in a marked decrease in bone strength and increase in cortical bone porosity. CRISPR/Cas9-mediated edits of *DAAM2* in osteoblast cell lines demonstrated a reduction in mineralization, compared to un-edited cells.

These newly discovered loci will empower future clinical and pharmacological research on osteoporosis, spanning from a better understanding of its genetic susceptibility to, potentially, biomarker discovery and drug targets. Moreover, to maximize the utility of these results to the community, all data are made freely available via web resources (see URLs). Below we summarize the key results from our investigations.

## Results

### GWAS for eBMD and Fracture

We selected 426,824 White-British individuals (55% female) for the eBMD GWAS from the UK Biobank full release (**Online Methods, Table S1** and **Figure S1**). We analyzed 13,737,936 autosomal and X-chromosomal SNPs for their association with eBMD. Although there was substantial inflation of the test statistics relative to the null for eBMD (λ_GC_ = 2.26, **Figure S2**), linkage disequilibrium (LD) score regression indicated that the majority of inflation was due to polygenicity rather than population stratification (LD score regression intercept = 1.06 [0.063], ratio = 0.017 [0.018]).

We identified 1,103 conditionally independent signals (423 novel) at a genome-wide significant threshold (P < 6.6×10^−9^ see **Online Methods**) mapping to 515 loci (301 novel) (**Table S2** and Figure 1). Of the conditionally independent lead SNPs at each locus, 4.6% were rare, having a minor allele frequency (MAF) ≤ 1%, whereas 9.3% were low-frequency (MAF ≤ 5% but > 1%) and 86.1% were common (MAF > 5%) (**Figure S3** shows the relationship between MAF and absolute effect size). The average absolute conditional effect sizes for these three categories of SNPs were 0.14, 0.04 and 0.02 standard deviations, respectively. The total variance explained by conditionally independent genome-wide significant lead SNPs for eBMD was 20.3%. When partitioning the variance explained by genome-wide significant lead SNPs into the three MAF categories, we found that rare variants explained 0.8% of the variance, whereas low-frequency and common variants explained 1.7% and 17.8% of the variance in eBMD, respectively. We found strong correlations between effect sizes for eBMD when compared to effect sizes from the interim release of UK Biobank data (r = 0.93, **Figure S4, Table S3**).

**Figure 1.**
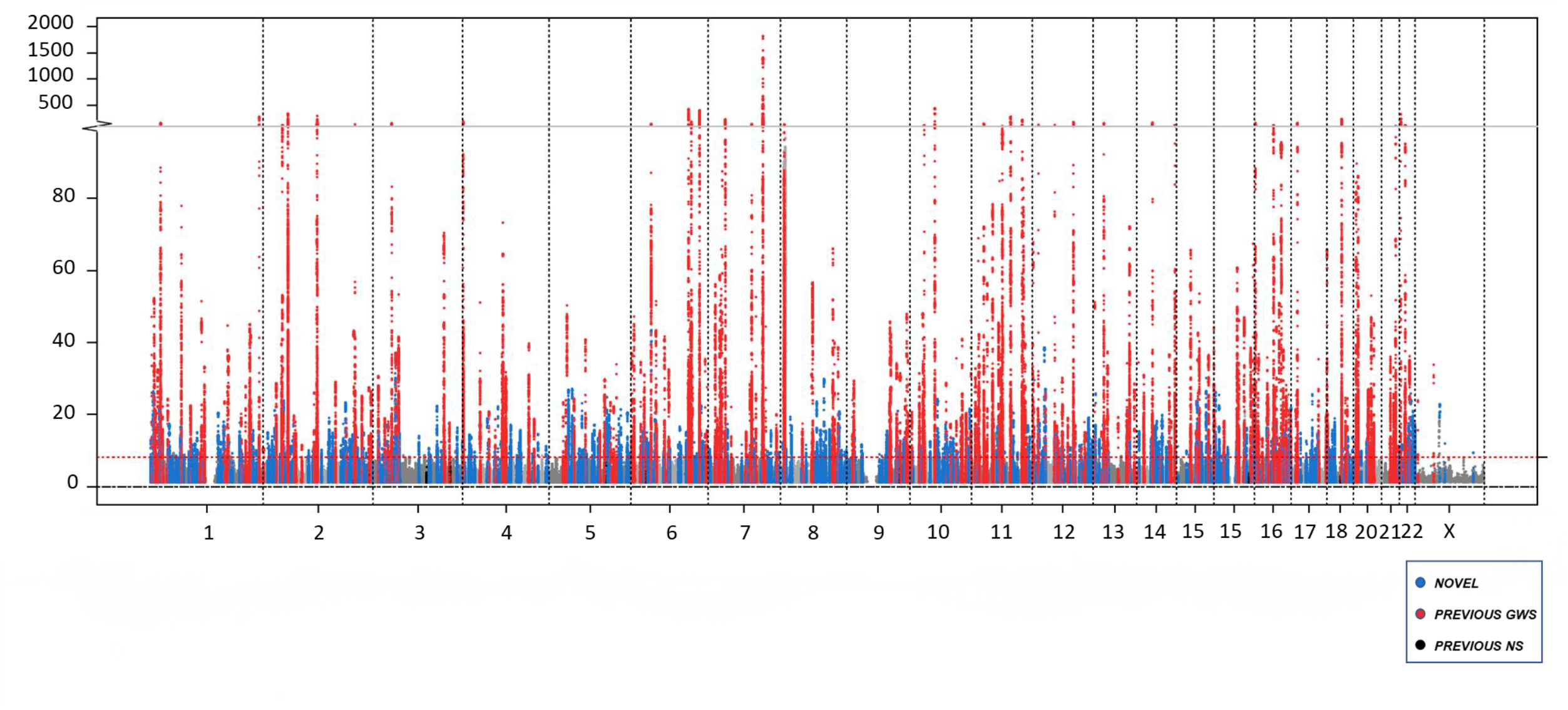
Manhattan plot of genome-wide association results for eBMD in the UK Biobank. The dashed red line denotes the threshold for declaring genome-wide significance (6.6×10^−9^). 1,103 conditionally independent SNPs at 515 loci passed the criteria for genome-wide significance. 301 novel loci (defined as > 1 Mbp from previously reported genome-wide significant BMD variants) reaching genome-wide significance are displayed in blue. Previously reported loci that reached genome-wide significance are displayed in red, and previously reported loci failing to reach genome-wide significance in our study are shown in black.

We identified 53,184 fracture cases (60% female) and 373,611 controls (54% female), totalling 426,795 individuals in UK Biobank (**Table S1**). We assessed 13,977,204 autosomal and X-chromosomal SNPs for their effects on fracture and identified 14 conditionally independent signals associated with fracture mapping to 13 loci (**Table S4 and Figure S5**). Once again, we observed inflation of the test statistics, (λ_GC_ = 1.15). However, this was also likely due to polygenicity, rather than population stratification (LD score regression intercept = 1.00 [0.008], ratio = 0.017 [0.038]). Conditionally independent genome-wide significant lead SNPs were tested for replication in a cohort of research participants from 23andMe, Inc., a personal genetics company (N = 367,900 cases and 363, 919 controls). All 14 SNPs showed strong evidence of replication (**Table S4**). All genome-wide significant fracture SNPs were also found to be genome-wide significant in their association with eBMD in the expected direction of effect (i.e. alleles lowering eBMD were related to higher risk of fracture). Further, there was a high correlation between the effect sizes of eBMD associated variants and their effects on fracture were highly negatively correlated (r = −0.77 [−0.79, −0.74], **Figure S4**).

### Sex Heterogeneity

To investigate whether the genetic aetiology of eBMD differed between the sexes, we performed tests of sex heterogeneity across the genome. We identified 45 variants at 7 loci that displayed strong evidence of a sex difference (P < 6.6×10^−9^, **Table S5**). Variants at two of these 7 loci did not reach genome-wide significance in males, females or the main eBMD GWAS, and were therefore not followed up further (**Figure S6** and **Table S5**). Of the five remaining loci (**Table S5**), we detected evidence of a sex difference at *FAM9B*, a known male-only eBMD associated locus that may mediate its effect on bone through both serum testosterone levels and estradiol levels in men.^13,14^ Alleles at this locus associated with increased testosterone levels were also associated with increased eBMD in males only. For the remaining loci, male-only effects were detected at *FKBP4* and *RNU6ATAC. FKBP4* codes for a tetratricopeptide repeat protein found in steroid receptor complexes that has been implicated in androgen receptor mediated signalling and function.^15^ Variants at the *LOC105370177* (upstream of the *OPG* gene) and *ABO* loci were associated with eBMD in both sexes, but were more strongly related in males. Finally, variants within *MCM8* were associated with eBMD in females only (**Table S6**). The same variants are known to be associated with onset of menopause^16^ in the predicted direction (i.e. alleles which increase age at menopause associate with increased eBMD). Interestingly, 164 loci that reached genome-wide significance in the main analysis showed evidence of sex-heterogeneity in effect size far above expectation (164 out of 1106 SNPs had P < 0.05, **Table S7**). LD score regression analyses suggested that the genetic architecture influencing male and female eBMD was largely shared but that there were some significant differences between the sexes (r_G_ = 0.91, SE=0.012, P < 0.001).^17^ The total number of genome-wide significant conditionally independent lead SNPs becomes 1,106 mapping to 518 loci when including our sex heterogeneity analyses, however, we focus on results from the main GWAS for the rest of our study.

### Coding Variants

Most genome-wide significant associations to date have arisen from non-coding variants, which has made the identification of causal genes difficult.^12^ Genetic association signals at coding variation can more directly highlight a potentially causal gene. We identified 1,237 coding variants, based on the Variant Effect Predictor^18^, meeting genome-wide levels of significance in their association with eBMD, prior to conditioning on other the lead SNPs in LD at each locus. This represents 1.0% of the total count of genome-wide significant variants (**Table S8**). The average absolute effect size for coding variants was 0.025 standard deviations (interquartile range: 0.014 – 0.027), which was approximately equal to the absolute effect size for genome-wide significant common variants. These coding variants do not necessarily directly implicate a gene but may reflect non-causal associations through linkage disequilibrium with other common non-coding causal variants.

### Fine-Mapping Associated Loci

In order to map SNPs to potentially causal genes, we first refined the set of associated SNPs at each locus to a smaller set using two statistical fine-mapping methods, GCTA-COJO^19^ and FINEMAP^20^. These methods identify sets of SNPs based on their conditional independence and posterior probability for causality, respectively. We generated such sets for each genome-wide significant autosomal locus by identifying conditionally independent lead SNPs, or those SNPs having a high posterior probability of causality, as determined by log_10_ Bayes factor > 3 (Figure 2a). Here we refer to the set of “fine-mapped SNPs” as those SNPs achieving either conditional independence or a high posterior probability for causality.

**Figure 2.**
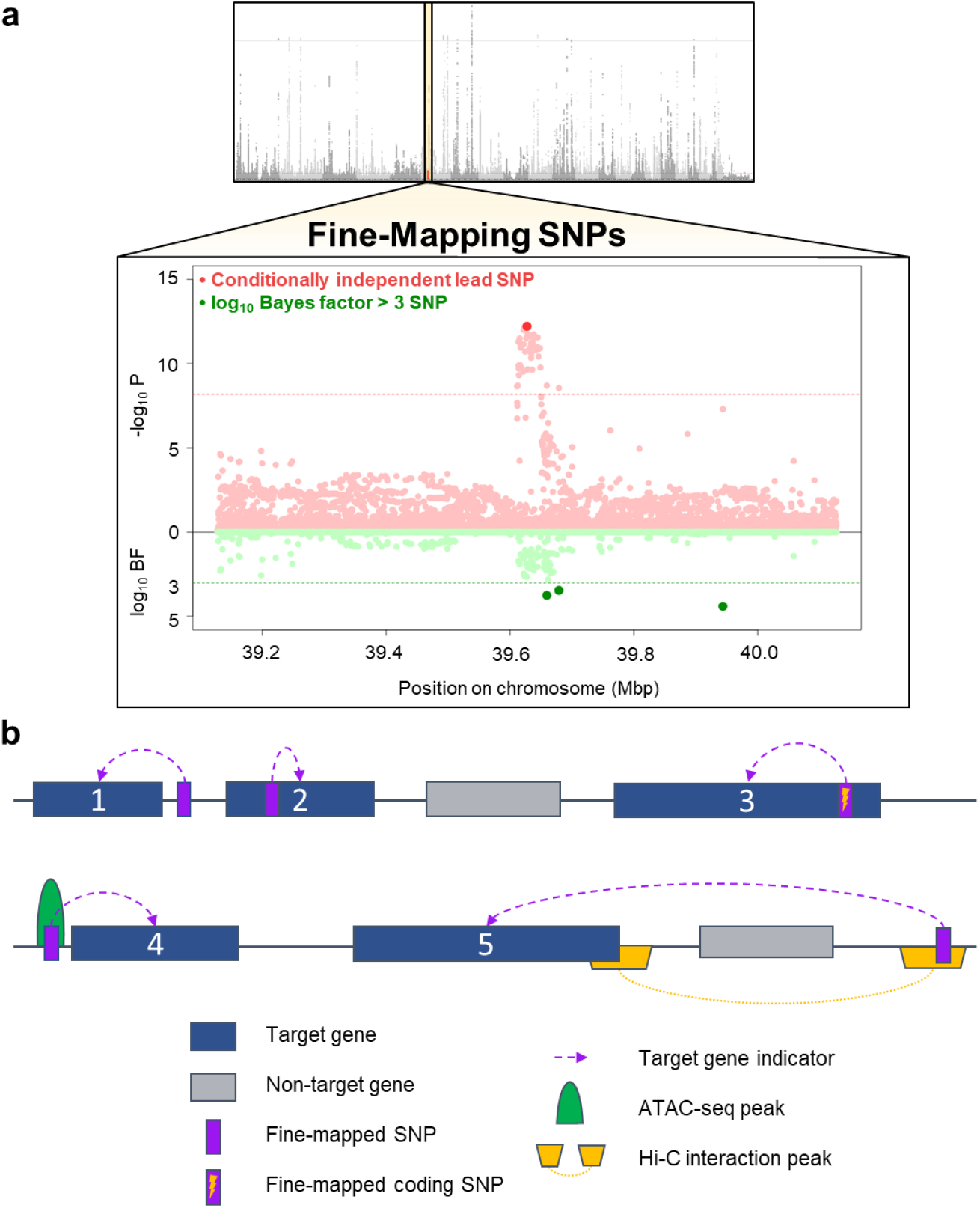
Fine-mapping SNPs and target gene selection diagram. **A**) For each 500 Mbp region around a conditionally independent lead SNP, we applied statistical fine-mapping to calculate log10 Bayes factors for each SNP as a measure of their posterior probability for causality. SNPs that were conditionally independent lead SNPs or that had log_10_ Bayes factors > 3 were considered our fine-mapped SNPs that we then used for target gene identification. **B**) Target Genes were identified if: 1) It was the gene closest to a fine-mapped SNP. 2) A fine-mapped SNP was in its gene body. 3) A fine-mapped SNP was coding. 4) The gene mapped closest to a fine-mapped SNP which resided in an SaOS-2 ATAC-seq peak. 5) A fine-mapped SNP was present in a Hi-C osteoblast or osteocyte promoter interaction peak, therefore being closer to a target gene in three-dimensions than linearly on the genome.

Prior to fine-mapping, we identified on average 235 genome-wide significant SNPs per locus. After this fine-mapping exercise, an average of two conditionally independent SNPs and five SNPs with a log_10_ Bayes factor > 3 remained per locus (**Tables S9** and **S10**). The number of fine-mapped SNPs per locus ranged between 1 to 81. As a sensitivity test, we also considered a more lenient inclusion criterion for inclusion of SNPs based on a log_10_ Bayes factor > 2, which resulted in a sharp increase in the average number of SNPs per locus to 27, which in total comprised 13,742 unique SNPs (**Table S11**).

### Comparing Fine-Mapped SNPs for Biological Activity

Given the large number of associated SNPs per locus, downstream analyses should focus on those SNPs most likely to have a biological function. We used accessible chromatin sites surveyed in a relevant cellular context as a proxy for biological activity. We generated ATAC-seq maps in the human osteosarcoma cell line SaOS-2. SaOS-2 cells possess osteoblastic features and can be fully differentiated into osteoblast-like cells. We also analyzed DNase I hypersensitive site (DHS) maps from human primary osteoblasts generated by the ENCODE project.^21^ Both ATAC-seq and DHS data were analyzed using a uniform mapping and peak-calling algorithm (**Online Methods**).

We then analyzed the fine-mapped SNPs for enrichment of these functional signatures relative to all SNPs in the 1 Mbp surrounding each genome-wide significant association locus. Fine-mapped SNPs, including the set of conditionally independent SNPs and SNPs with log_10_ Bayes factors > 3, were strongly enriched for both missense variants in protein coding regions and osteoblast accessible chromatin sites (Figure 3a). As the log_10_ Bayes factor threshold increased, fold-enrichment increased as well (Figure 3b). This indicates that the fine-mapped set of SNPs is highly enriched for genomic signatures of function, which can inform the choice of statistical cut-off for selection of SNPs for follow-up functional studies.

**Figure 3.**
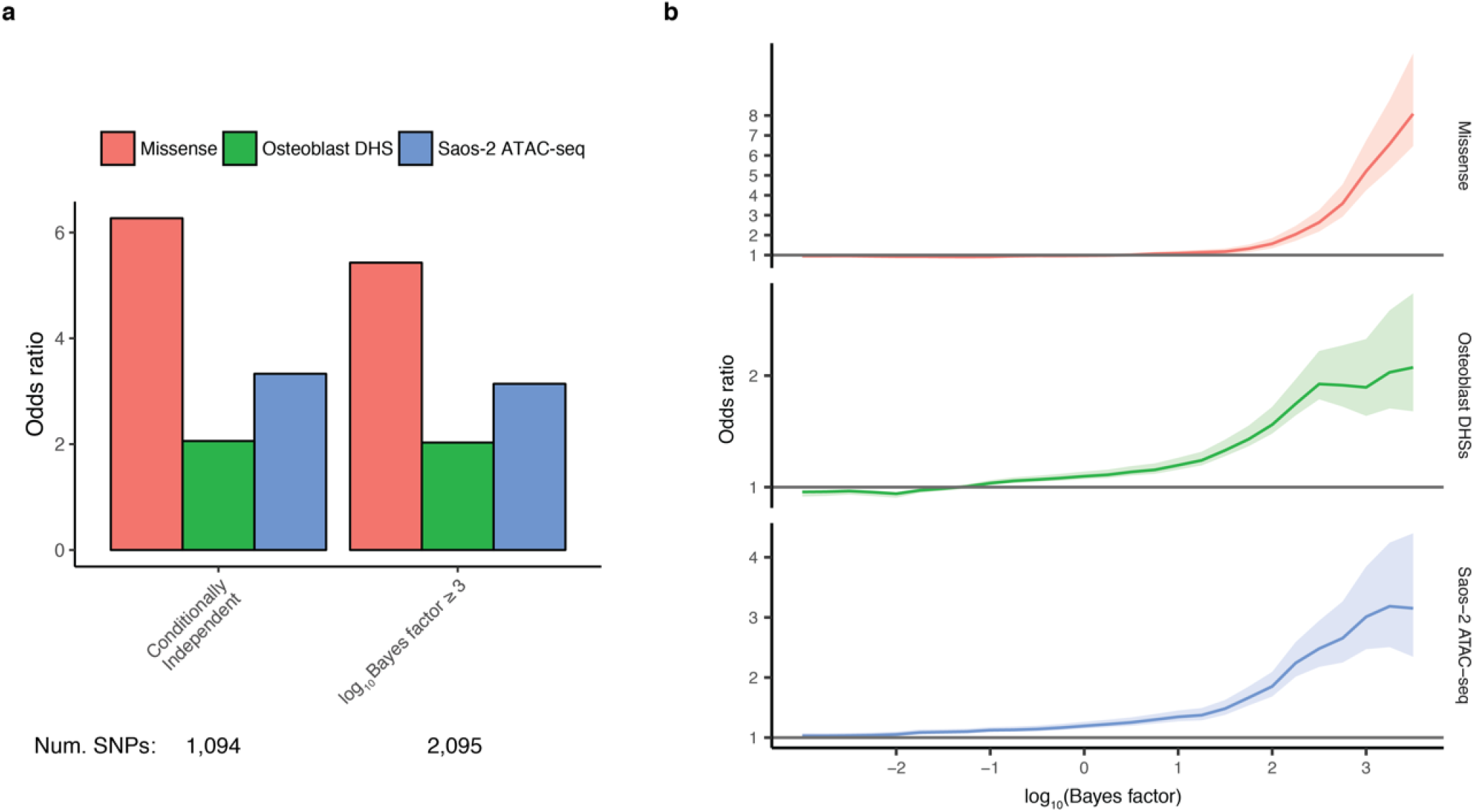
SNPs at genome-wide significant loci are enriched for osteoblast open chromatin sites. **A**) Odds ratio for missense, osteoblast DHSs and SaOS-2 ATAC-seq peaks for SNPs that are conditionally independent or achieving a log10 Bayes factor > 3. Note the log10 Bayes factor > 3 set contains nearly twice the number of SNPs. **B**) Ranking SNPs by log10 Bayes factor (x-axis) shows increasing enrichment of missense SNPs and of SNPs at accessible chromatin sites.

### Mapping Fine-Mapped SNPs to Target Genes & Enrichment for Positive Control Genes

Human genetic associations have rarely been translated to improved clinical care, primarily because causal genes at associated loci have not been indisputably identified. We therefore sought to test which genomic features link associated SNPs to genes known to influence bone biology in humans. We identified a set of proteins whose perturbation through pharmacotherapy^2^ or Mendelian disease leads to changes in bone density or strength. Mendelian disease genes were defined as monogenic disorders characterized with altered bone mass or abnormal skeletal mineralization, osteolysis and/or skeletal fragility or osteogenesis imperfecta (**Table S12**) and constitute an informative human knockout resource.^22^ We considered such proteins to be products of “positive control” genes influencing bone density and likely critical to bone biology.

Next, we investigated which genomic features linked the fine-mapped set of SNPs to positive control genes for bone density. We tested whether positive control genes were enriched among six types of genomic characteristics that can link a SNP to a gene: 1) Genes that were most proximal to the fine-mapped set SNPs; 2) Genes that contained fine-mapped SNPs overlapping their gene bodies; 3) Genes containing fine-mapped SNPs that are coding variants; 4) Genes identified to be in 3D contact with fine-mapped sets in human osteoblasts or osteocytes through high-throughput chromatin conformation capture (Hi-C) experiments; 5) The closest gene to fine-mapped SNPs, which also mapped to ATAC-seq peaks in human osteoblast SaOS-2 cell lines; and 6) Those genes within 100 kbp of fine-mapped SNPs (Figure 2b emphasizes the target gene selection and Figure 4 details this entire pipeline). Coding annotations, ATAC-seq peaks, and Hi-C interaction peaks were not combined but kept separate to enable different sources of data to provide converging and confirmatory evidence. Distance from a fine-mapped SNP to a gene was considering the closer of the 3’ and 5’ ends, not the transcription start site. We named these genes “Target Genes” and tested which of the above 6 methods to define Target Genes was most strongly enriched for positive control genes.

**Figure 4.**
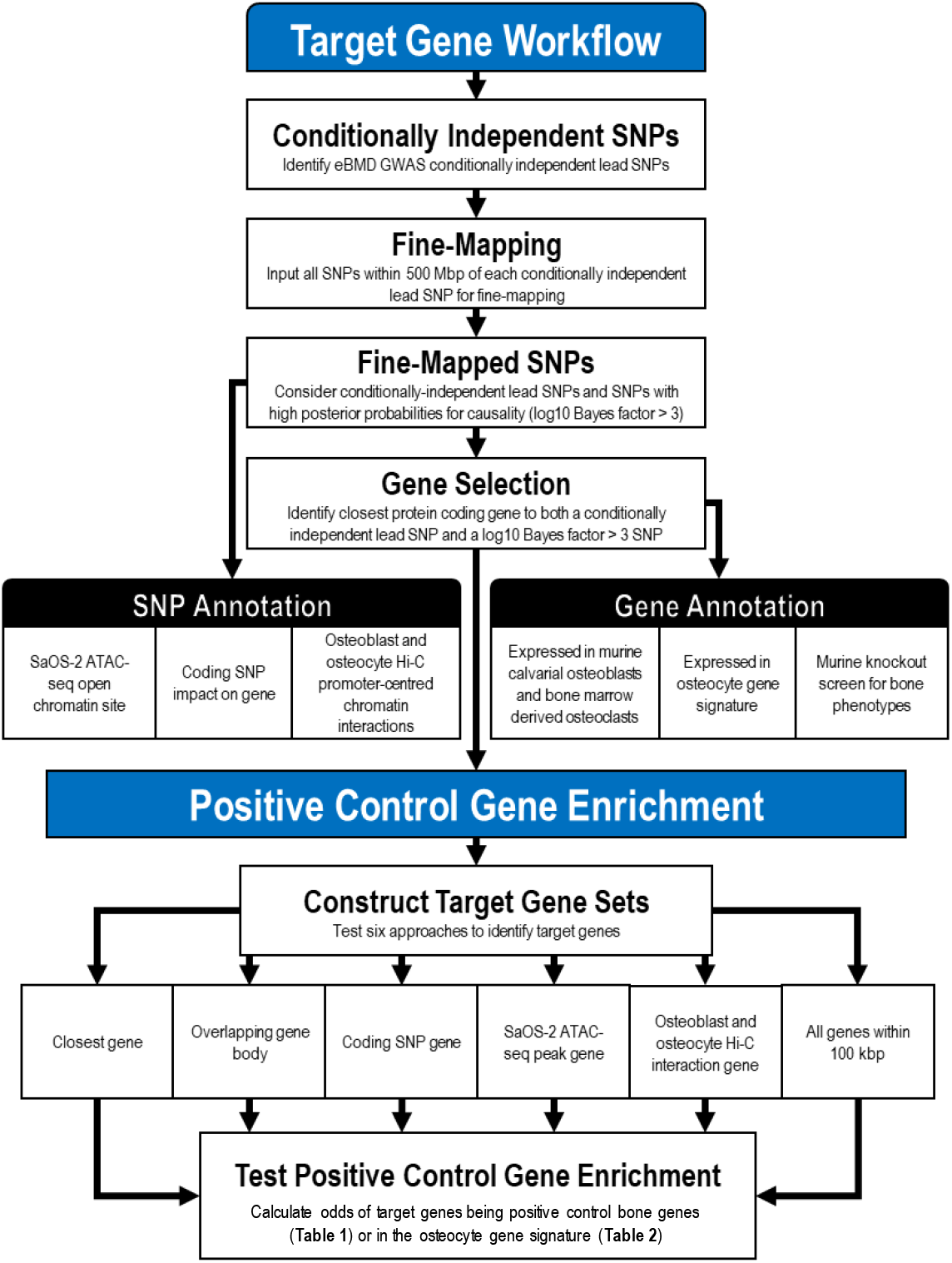
Target Gene Identification Workflow.

The set of Target Genes that were most strongly enriched for positive control genes, arose from genes targeted by SNPs that were conditionally independent and by SNPs identified to be plausibly causal with a log_10_ Bayes factor > 3 (Table 1 **and Table S13**). This set of Target Genes featured 556 genes total, approximately one gene per locus. All six different methods for linking these fine-mapped set of SNPs to Target Genes yielded strong enrichment for positive control genes. The odds ratios ranged from 5.1 (95% CI: 3.0-8.6, P = 10^−11^) for Target Genes within 100 kbp of the fine-mapped SNPs to an odds ratio of 58.5 (95% CI: 26.4-129.31, P = 10^−75^) for Target Genes closest to fine-mapped SNPs that were in an osteoblast-derived ATAC-seq peak (Table 1). In addition, we used FUMA^23^ to assess which pathways from the WikiPathways^24^ database were identified by the set of Target Genes most strongly enriched for positive control genes. We observed that well known pathways such as Wnt signalling, endochondral ossification, osteoclast and osteoblast signalling, as well as novel pathways were highlighted by this approach (**Figure S7**).

**Table 1.** Target gene identification methods enrichment for 57 positive control genes. No positive control genes were identified by osteocyte Hi-C interactions therefore we did not calculate its enrichment. Distance to gene was determined using 3’ and 5’ ends, instead of the transcription start site.

| *Target Gene Set* | *OR (95% CI)* | *P* |
| --- | --- | --- |
| *SaOS-2 ATAC-seq Peak Gene* | 58.5 (26.4 – 129.3) | $1.3 \times 10^{-75}$ |
| *Coding SNP Gene* | 41.8 (14.3 – 121.6) | $1.0 \times 10^{-30}$ |
| *Osteoblast Hi-C Interaction Gene* | 21.1 (6.4 – 69.6) | $7.8 \times 10^{-13}$ |
| *Closest Gene* | 12.9 (7.1 – 23.4) | $1.8 \times 10^{-27}$ |
| *Overlapping Gene Body* | 11.2 (5.2 – 23.8) | $3.4 \times 10^{-15}$ |
| *All Genes Within 100 kbp* | 6.8 (3.9 – 11.7) | $2.1 \times 10^{-15}$ |
| *Osteocyte Hi-C Interaction Gene* | NA | NA |

These results suggest that our Target Gene identification methods lead to strong enrichment for positive control genes known to be central to bone biology. Such methods may help to prioritize genes at associated loci for functional testing, which are more likely to influence bone biology and therefore, have clinical relevance. The full list of mapped Target Genes and the method through which they were identified is presented in **Table S14**.

### Mapping Fine-Mapped SNPs to Osteocyte-Signature Genes

An alternative method to assess the biological plausibility of Target Genes is to test whether their expression is enriched in bone cells. Osteocytes are the most abundant cell type in bone and are key regulators of bone mass, bone formation and bone resorption.^25^ We therefore assessed the transcriptome of primary murine osteocytes derived from three bone types *in vivo.^26^* Genes enriched for expression in osteocytes and expressed in all bone types defined an osteocyte transcriptome signature.^26^ We then tested which of the methods used to identify eBMD Target Genes resulted in the greatest enrichment for osteocyte-signature genes.

Again, we found that Target Genes were strongly enriched for osteocyte signature genes, with odds ratios for enrichment ranging from 2.1 (95% CI: 1.7-2.5, P = 2×10^−17^) for Target Genes within 100 kbp of the fine mapped set of SNPs, to 7.4 (95% CI: 3.8-14.5, P = 5×10^−12^) for Target Genes mapped through fine-mapped coding SNPs (Table 2 **and Table S15 and S16**). This again suggests our methods result in enrichment for biologically relevant genes.

**Table 2.**
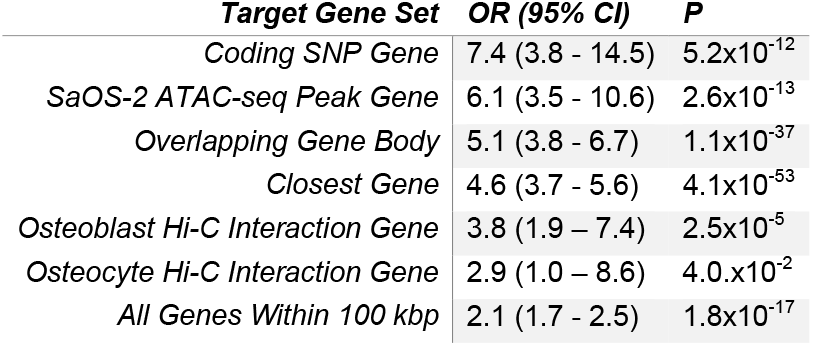
Target gene identification methods enrichment for 1,240 osteocyte signature genes. Distance to gene was determined using 3’ and 5’ ends, instead of the transcription start site.

### A Large-Scale High Throughput Murine Knockout Screening Program

The Origins of Bone and Cartilage Disease (OBCD) program (www.boneandcartilage.com) is determining 19 structural and functional parameters in all unselected knockout mouse lines generated at the Wellcome Trust Sanger Institute for the IKMC and IMPC. These parameters evaluate bone mineral content (BMC), 3D trabecular and cortical bone structure, bone mineralization and femoral and vertebral bone strength. To date, the OBCD program has included the analysis of 126 knockout lines with mutations of Target Genes (**Table S17**). Outlier phenotypes were defined as structural or strength parameters > 2 standard deviations away from the reference mean, determined from over 300 age-matched, sex-matched and genetically identical C57BL/6N wild-type controls (**Online Methods**). We investigated whether deletion of these 126 Target Genes resulted in enrichment of outlier skeletal phenotypes. Outlier cortical and trabecular bone phenotypes were more frequent in mice with disruptions of the 126 Target Genes compared against 526 unselected knockout lines (**Tables S17 and S18**, OR 3.2 [95% CI: 1.9-5.6], P < 0.0001). Therefore, enrichment of abnormal skeletal phenotypes in mice with disruption of Target Genes provides clear functional validation that our fine-mapping approach identifies critical and biologically-relevant skeletal genes. Our fine-mapping *in vivo* and *in vitro* data converged to identify *DAAM2* as a highly credible and novel osteoporosis gene, therefore we undertook detailed analyses of mice with a hypomorphic *Daam2* allele to illustrate the potential of this approach.

### In-Depth Characterization of DAAM2

Numerous lines of evidence identified *DAAM2* as an important gene for further functional investigation. First, a conditionally independent lead SNP, rs2504101, mapped directly to *DAAM2* (P_conditional_ = 4.3 x 10^−10^) and second, fine-mapping revealed two coding missense variants with high posterior probabilities for causality, rs201229313 in its 19^th^ exon (log_10_ BF = 3.7), and rs61748650 in its 21^st^ exon (log_10_ BF = 2.5). Third, a rare variant, rs772843886, near *DAAM2* was suggestively associated with risk of fracture (P = 2×10^−3^). Fourth, the *Daam2^tm1a/tm1a^* mouse was identified to have an outlier skeletal phenotype in our rapid throughput murine knockout screening program (**Table S17**). Fifth, although *DAAM2* has not previously been implicated in osteoporosis, it has been predicted to have a role in canonical Wnt signaling.^27,28^

To investigate the role of *DAAM2* in bone biology, we first tested its expression in bone cells. We performed RNA-seq and ATAC-seq experiments in four different human osteoblast cell lines and found it was expressed in all cell lines (**Online Methods, Figure S8**). Staining experiments in the SaOS-2 cell line revealed DAAM2 localized specifically in the cell nuclei (**Figures S9 and S10**). This functional evidence from human bone cells also led us to characterize *Daam2* in mouse bone cells. *Daam2* was identified as an osteocyte signature gene (**Table S16**) and was expressed in mouse calvarial osteoblasts and bone marrow-derived osteoclasts (**Table S19**).

Next using CRISPR/Cas9, we tested the effect on bone mineralization of double-stranded breaks (DSBs) in the second exon of *DAAM2* in SaOS-2 osteoblast cell lines (**Online Methods**). We found that after 14 days of treatment with osteogenic factors, control cells transfected with the intact plasmid, but not undergoing an DSB of the *DAAM2* gene, had a 9-fold increase in mineralization. After the introduction of a DSB in the second exon of *DAAM2*, induced mineralization was severely impaired (Figure 5). These CRISPR/Cas9-based findings suggest that DAAM2 influences mineralization capacity in human osteoblasts.

**Figure 5.**
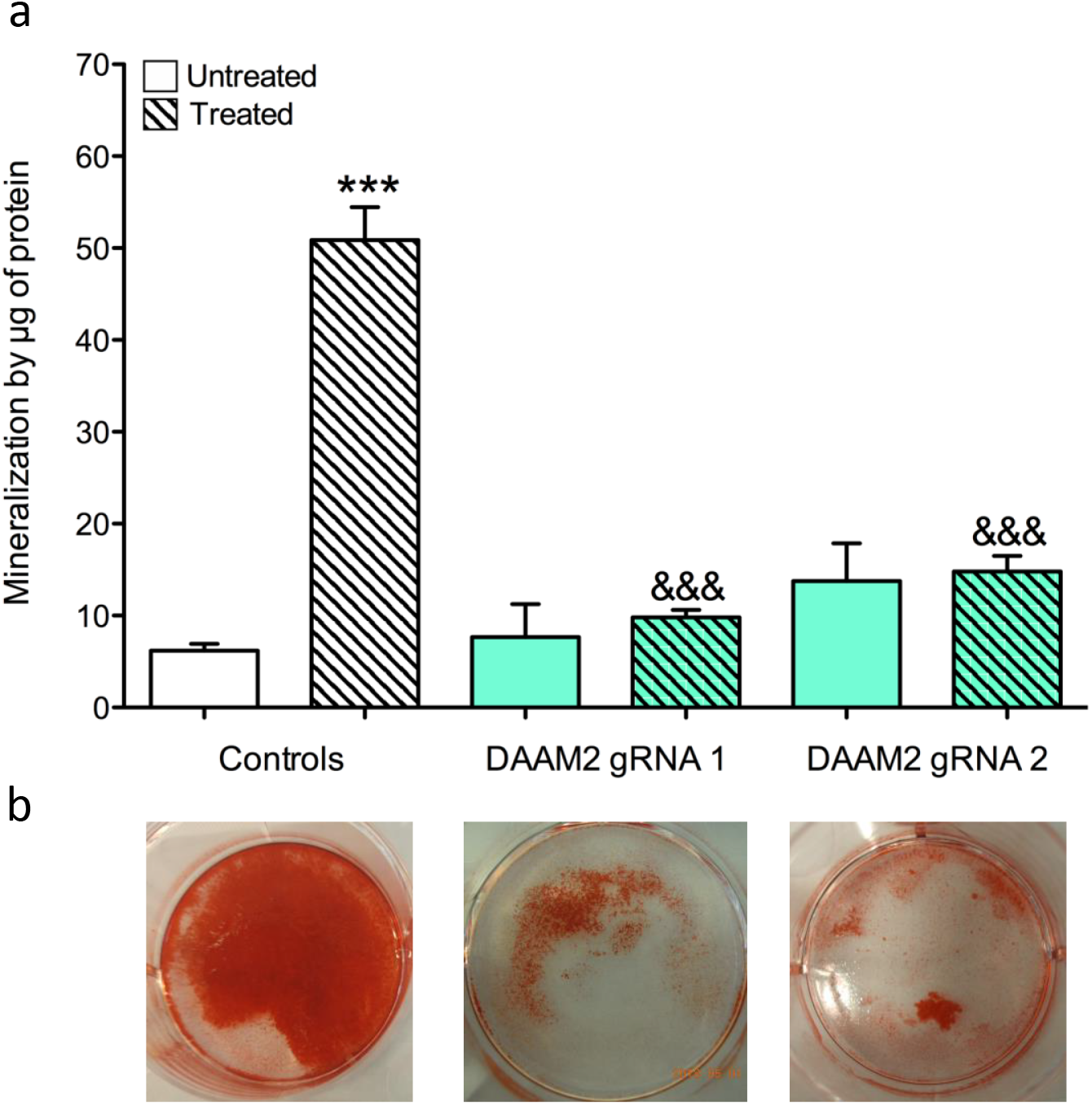
Reduction in DAAM2 protein resulted in decreased mineralization in SaOS-2 cells. Mineralization quantification in control cells and *DAAM2* exon 2 double-stranded break (DSB) induced cells in either the presence of osteogenic factors (treated) or absence (untreated). Bars in (**a**) represent the mean of six independent experiments ± SEM from Alizarin red staining in (**b**) to quantify mineralization. *** P < 0.001 compared to untreated control cells and &&& P < 0.001 compared to treated control cells determined by one-way Anova and a Bonferroni post-hoc test.

We next analyzed the skeletal phenotypes of *Daam2^tm1a/tm1a^, Daam2^+/tm1a^* and wild-type littermate mice in detail. Adult male *Daam2^tm1a/tm1a^* mice had reduced femur and vertebral bone mineral content (BMC), while male *Daam2+^tm1a^* and female *Daam2^tm1a/tm1a^* mice also had reduced vertebral BMC. These changes were accompanied by a small reduction in femur length in *Daam2^tm1a/tm1a^* mice (males, 2.7%; females, 3.5%). Despite otherwise normal trabecular and cortical bone structural parameters, cortical porosity was increased in both male and female *Daam2^tm1a/tm1a^* mice (**Figure S11**).

Consistent with their increased cortical porosity, *Daam2^tm1a/tm1a^* mice had markedly reduced bone strength (Figure 6) even though all other cortical bone parameters, including BMD, were normal (**Figure S11**). Bone composition and structure were thus investigated in *Daam2^tm1a/tm1a^* mice by comparing *Daam2^tm1atm1a^* mineralization and biomechanical parameters with values predicted by linear regression analysis of over 300 wild-type age, sex and genetic background matched wild-type controls. Measures of bone composition and structure in *Daam2^tm1a/tm1a^* mice were reduced compared to wild-type mice, and vertebral stiffness was > 2 standard deviations below that predicted even after accounting for reduced BMC (Figure 6c **and Table S20**). To investigate the role of *Daam2* on bone turnover, we measured markers of bone resorption (TRAP) and formation (P1NP) in 10-week-old *Daam2^tm1atm1a^* and *Daam2+^tm1a^* mice, and these did not differ from wild-type (**Figure S12**). Furthermore, primary cultures of bone marrow mononuclear cells from *Daam2^tm1a/tm1a^* mice showed no difference in osteoclastogenesis, and primary osteoblast mineralization was also similar to wild-type (**Figure S12**).

**Figure 6.**
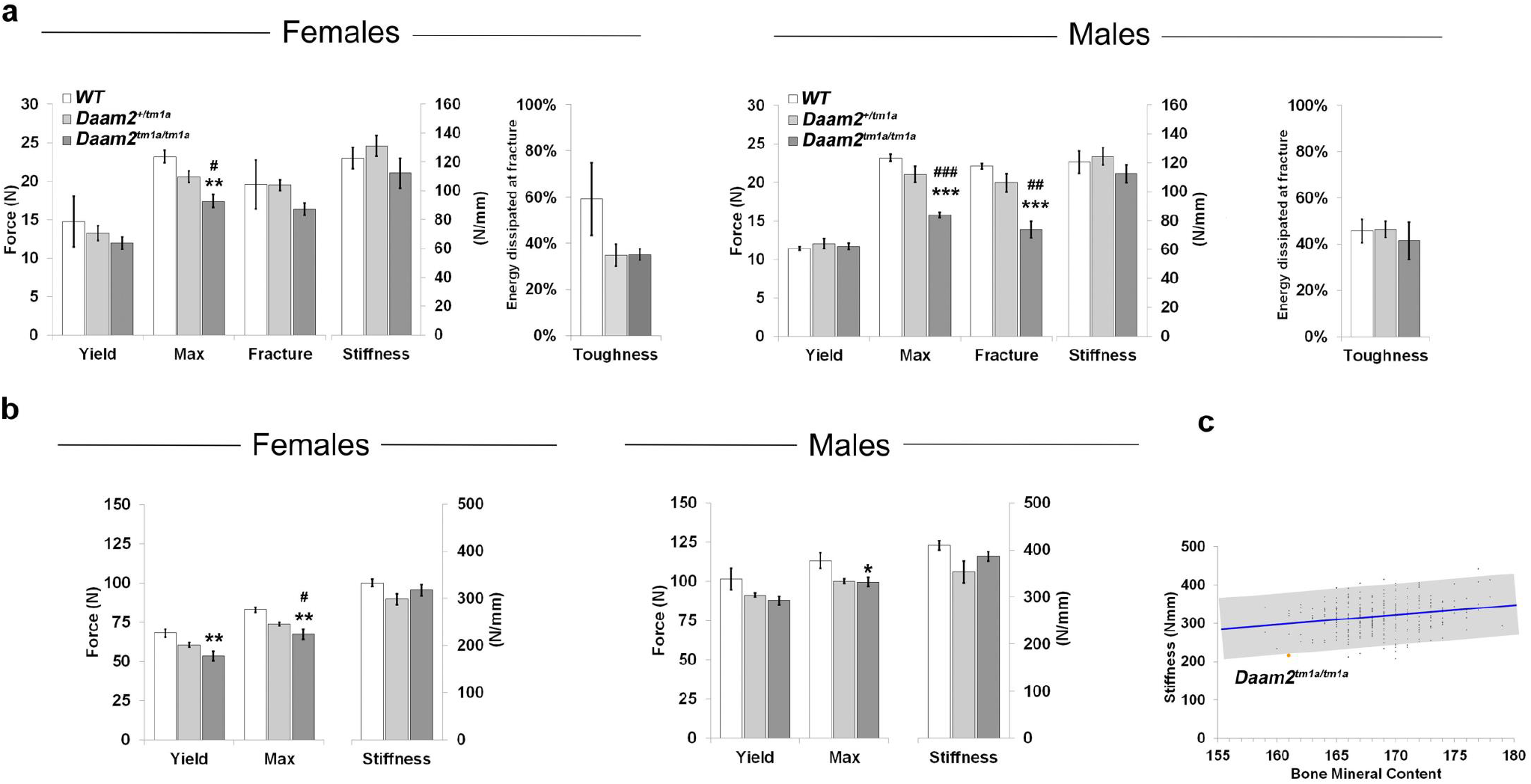
Biomechanical Analyses of mice with Daam2 knockdown. **A) Femur biomechanical analysis.** Destructive 3-point bend testing (Instron 5543 load frame) of femurs from WT (N_Female_ = 3, N_Male_ = 4), *Daam2^+/tm1a^* (N_Female_ = 6, N_Male_ = 4), *Daam2^tm1a/tm1a^* (N_Female_ = 5, N_Male_ = 9) mice. Graphs showing yield load, maximum load, fracture load, stiffness (gradient of the linear elastic phase) and toughness (energy dissipated prior to fracture). Data are shown as mean ± SEM; ANOVA and Tukey’s post hoc test; (i) *Daam2^+/tm1a^* vs WT and *Daam2^tm1a/tm1a^* vs WT, **P<0.01; ***P<0.001 and (ii) *Daam2^+/tm1a^* vs *Daam2^tm1a/tm1a^*, #P<0.05; ##P<0.01; ###P<0.001. **B) Vertebra biomechanical analyses**. Destructive compression testing (Instron 5543 load frame) of caudal vertebrae from WT (NFemale = 3, NMale = 4), *Daam2^+/tm1a^* (NFemale = 6, NMale = 4), *Daam2^tm1a/tm1a^* (N_Female_ = 5, N_Male_ = 9) mice. Graphs showing yield load, maximum load, and stiffness. Data are shown as mean ± SEM; ANOVA and Tukey’s post hoc test; (i) *Daam2^tm1a/tm1a^* vs WT, *P<0.05 and **P<0.01 and (ii) *Daam2^+/tm1a^* vs *Daam2^tm1a/tm1a^*, #P<0.05. Females are on left and males on right. **C) Bone composition and structure analysis from rapid throughput screening murine knockouts**. The graph demonstrates the physiological relationship between bone mineral content and stiffness in caudal vertebrae from P112 female WT mice (N = 320). The blue line shows the linear regression (P = 0.0001) and the grey box indicates ± 2SD. The mean value for female *Daam2^tm1a/tm1a^* (N = 2 from initial OBCD screen) mice is shown in orange (−2.14 SD).

Male *Daam2^tm1a/tm1a^* mice had decreased mineral content per unit matrix protein and increased carbonate substitution (**Figure S13**). This decrease in mineral to matrix ratio explains the overall decrease in bone mineral content observed in the absence of a decrease in cortical bone size. While bone size and geometry play a major role in controlling bone strength, decreases in mineral to matrix ratio are associated with decreased bone stiffness and decreased bending moment.^29^ These decreases likely contributed to the poor bone composition and structure observed in the *Daam2^tm1a/tm1a^* mice.

Taken together, these data suggest the decreased bone strength in *Daam2^tm1a/tm1a^* mice is not simply a result of abnormal bone turnover, but also a consequence of increased porosity and impaired bone composition and structure. If DAAM2 proves to be a tractable drug target, such an agent would represent a complementary therapeutic strategy for prevention and treatment of osteoporosis and fragility fracture.

### Additional Novel Candidate Bone Genes

While *DAAM2* represents the detailed validation of a novel Target Gene and the rapid-throughput knockout mouse skeletal phenotyping pipeline, we also highlight five additional eBMD Target Genes that result in contrasting abnormalities of bone structure and strength when deleted in mice, thus emphasising their functional role in skeletal physiology and importance for further study.

*CBX1* encodes Chromobox 1, a highly conserved non-histone member of the heterochromatin protein family that mediates gene silencing but has no reported role in the skeleton^30^. Homozygous deletion of *Cbx1* resulted in embryonic lethality whereas adult heterozygous mice had increased bone mineral content and trabecular thickness resulting in increased stiffness and strength (**Table S17, Figure S14**). *CBX1* was identified by five SNPs with log10 BFs > 2 mapping directly to its gene body (**Table S11**) and rs208016 (70 kbp upstream) suggested an association with fracture (P = 1.5×10^−5^).

*WAC* encodes WW Domain Containing Adaptor with Coiled-Coil, a protein of unknown function that is associated with global developmental delay and dysmorphic features in Desanto-Shinawi syndrome^31^. Homozygous deletion of *Wac* resulted in prenatal lethality whereas adult heterozygous mice had increased bone length, mass and strength (**Table S17, Figure S15**). Seven fine-mapped SNPs mapped proximally or directly to *WAC* (**Table S11**), with two fine-mapped SNPs, rs17686203 (log10 BF = 3.1) and rs61848479 (log10 BF = 3.9) mapping to *WAC* promoter Hi-C interaction peaks in primary human osteoblasts, and for the latter SNP in primary human osteocytes (**Table S14**). We also identified rs17753457 (60 kbp downstream) that had a suggestive association with fracture (P = 4.3×10^−5^).

*DSCC1* encodes DNA Replication and Sister Chromatid Cohesion 1, a component of an alternative replication factor that facilitates binding of proliferating cell nuclear antigen to DNA during S phase but has no known role in bone^32^. Homozygous knockout mice had reduced viability and adult *Dscc1^+/−^* heterozygotes had increased bone mineral content and strength (**Table S17, Figure S16**). *DSCC1* was identified by rs62526622 (log10 BF = 2.0) mapping to an intronic *DSCC1* Hi-C promoter interaction peak in primary human osteoblasts. rs546691328 (180 kbp downstream) was also found to have a suggestive association with fracture (P = 2.9×10^−4^).

*RGCC* encodes Regulator of Cell Cycle, a p53 Target Gene that interacts with polo-like kinase 1, which regulates cell proliferation and apoptosis but has no documented role in the skeleton^33^. Nevertheless, *Rgcc^−/−^* knockout mice displayed increased bone mineral content and strength (**Table S17, Figure S17**). *RGCC* was identified by rs145922919 (log10 BF = 3.3) mapping approximately 30 kbp upstream of *RGCC* to a Hi-C promoter interaction peak in primary human osteoblasts and rs545753481 (32 kbp upstream) also had a suggestive association with fracture (P = 3.4×10^−3^).

*YWHAE* encodes Tyrosine 3-Monooxygenase/Tryptophan 5-Monooxygenase Activation Protein, Epsilon Isoform, a pro-inflammatory cytokine that mediates signal transduction by binding to phosphoserine-containing proteins. YWHAE (14-3-3ε) binds to aminopeptidase N (CD13) to regulate chondrocyte homeostasis and has been implicated as a novel therapeutic target in osteoarthritis^34^. Rare *YWHAE* deletions have been reported in Miller-Dieker Lissencephaly syndrome which includes craniofacial abnormalities and growth retardation together with diverse neurodevelopmental abnormalities^35^. Consistent with this, homozygous deletion of *Ywhae* resulted in reduced bone length, and increased bone mass and mineral content resulting in brittle bones (**Table S17, Figure S18**). *YWHAE* was identified in our target gene approach by 22 SNPs with log10 BFs > 2 (**Table S11**) all mapping directly to *YWHAE* introns and an additional SNP, rs181451348 (1 kbp downstream) showed suggestive association with fracture (P = 7.1×10^−5^).

*CBX1, DSCC1, RGCC, WAC*, and *YWHAE* represent strong candidates for further in-depth functional characterization as we have performed for *DAAM2*. Bone composition and structure screens identified *WAC* and *DSCC1* as femur outliers due to *Wac^+/−^* and *Dscc1^+/−^* knockout mice being at least two standard deviations from the expected range (**Figure S19**). Our data also support functional experiments in human cells as all five genes were expressed in all four human osteoblast cell lines we profiled with RNA-seq and ATAC-seq (**Online Methods**), except for *RGCC* which was highly expressed in SaOS-2 with low expression levels in U2OS, MG63, and HOS, three other human osteoblast cell lines for which we generated RNA-seq data (**Online Methods**). In addition, we observed suggestive association at each locus with fracture (**Table S21**), further supporting evidence for these five genes having roles in human bone biology.

## Discussion

In this, the most comprehensive human and murine study on the genetic determinants of bone density and fracture performed to date, we have identified a total of 518 genome-wide significant loci, of which 301 are novel and together explain 20% of the total variance in eBMD. In a GWAS meta-analysis of up to 1.2 million individuals, 13 fracture loci were identified, all of which were also associated with eBMD. Taking advantage of the polygenicity of eBMD, we demonstrated strong biological enrichment for fine-mapped SNPs in bone cell open chromatin. Using fine-mapped SNPs we found that Target Genes were strongly enriched for genes that are known to play central roles in bone biology through Mendelian genetics, or as targets for clinically-validated osteoporosis therapies. High throughput skeletal phenotyping of mice with deletions of 126 Target Genes revealed enrichment of outlier skeletal phenotypes compared to analysis of 526 unselected knockout lines. Last, we identified DAAM2 as a protein with critical effects on bone strength, porosity and composition. These findings will enable on-going and future studies to better understand the genomic characteristics that link fine-mapped SNPs to sets of genes enriched for causal proteins. Further, this comprehensive study of the genetic variants associated with osteoporosis will provide opportunities for biomarker and drug development

The polygenicity of eBMD is striking. Few traits and diseases currently have hundreds of loci associated at genome-wide levels of significance.^12,36^ This has led to a large proportion of total variance in eBMD being explained by now known genetic determinants, which will facilitate future exploration of bone biology and enable drug development for osteoporosis.^37^, Yet, despite the large number of genetic and biological inputs into eBMD determination, pharmacological perturbation of even only one protein identified in our GWAS can have clinically relevant effects. For example, RANKL inhibition has been shown to increase bone density by up to 21% after ten years of therapy.^38^ Interestingly, the genetic variants near RANKL have small effects on eBMD. Thus, despite the small effect sizes for most identified variants, these do not necessarily reflect the effect sizes to be anticipated by pharmacological manipulation of the protein. This is because common genetic variants tend to have small effects on protein function, whereas pharmacotherapies tend to have large effects on protein function. Consequently, the dose-response curve describing the effect of small and large genetic perturbations on eBMD is needed to decide which proteins to target for drug development.^12^

Polygenicity has also improved our statistical power to validate linking an associated locus with a potentially causal gene. We found that fine-mapped sets of SNPs were able to identify Target Genes that were strongly enriched for positive control genes—particularly when the approach implemented relatively simple strategies, such as the nearest gene, or the gene nearest a fine-mapped SNP in cell-relevant open chromatin. We also observed that fine-mapped SNPs were often in 3D contact with Target Genes in human osteoblasts and osteocytes. These rich data, surveying many genomic landscape features provide guidance for investigators attempting to identify causal genes from GWAS-associated SNPs and all human genetic and murine results are available for download (see URLs).

The marked reduction in bone strength in *Daam2^tm1a/tm1a^* mice, despite minimal changes in bone morphology and mineral content, indicates that *Daam2^tm1a/tm1a^* mice have abnormal bone composition and structure, which can be explained in part by increased cortical porosity. Further, CRISPR/Cas9-mediated knockouts of *DAAM2* in osteoblast cells lines resulted in a marked reduction in inducible mineralization. Few such genes have been identified and further investigations will be required to determine whether *DAAM2* represents a tractable drug target in humans. Nevertheless, previous studies have suggested that *DAAM2* indirectly regulates canonical Wnt signalling across several developmental processes.^27,28^ Using different sources of data to identify *DAAM2*, allowed for greater confidence in results. While each type of data has its own biases, these biases are partially orthogonal, and consequently, concordant evidence from different sources of data increases the quality of the evidence, an approach known as triangulation.^39^

Our GWAS for fracture risk identified 13 loci associated with this common disease. All these loci have been associated with BMD and/or eBMD, highlighting the importance of BMD as a determinant of fracture risk, at least in the age range assessed within the UK Biobank. While BMD-independent loci for fracture likely exist, these were not identified despite a well-powered study. This suggests that screening for fracture drug targets should also include understanding the effect of the protein on BMD.

Our study has important limitations. First, we have measured eBMD, rather than DXA-derived BMD, which is typically measured in the clinic. Nonetheless, beyond their phenotypic correlation, these two traits also demonstrate high genetic concordance in terms of their genome-wide significant loci, suggesting that the biological properties that underpin these two traits are similar. Importantly, however, eBMD is a strong predictor of fracture risk in its own right, and contributes to risk assessment over and above DXA-derived BMD at the hip.^40^ While our target gene approach has identified a set of candidate genes enriched for genes with known effects on bone density, it is important to note that there is no gold-standard set of genes known to influence BMD. While our rapid throughput mouse knockout program is on-going and will investigate many of the Target Genes implicated by our study, further efforts will be required to functionally validate (or exclude) these genes in bone biology. Our target gene approach did not include human gene expression quantitative trait loci (eQTL) data. This is because the largest available eQTL experiments for human osteoblasts involve only 95 individuals,^41^ and larger sample sizes with RNA-sequencing data will be required to properly investigate our method of linking fine-mapped sets of SNPs to genes. Finally, our program was limited to individuals of White-British genetic ethnicity and the effect of most of the genome-wide significant SNPs in other populations remains to be assessed. It is likely that on-going studies in non-British ancestries will address this question.

In summary, we have generated an atlas of human and murine genetic influences on osteoporosis. This comprehensive study has more fully described the genetic architecture of eBMD and fracture and has identified a set of Target Genes strongly enriched for genes with known roles in bone biology. We used human genetics, functional genomics, animal models and genome editing to demonstrate the relevance of this approach, formally known as triangulation^39^, by identifying *DAAM2*. Disruption of *DAAM2* in mice leads to an increase in cortical porosity and marked reductions in bone composition and strength, and in human osteoblasts leads to a decrease in mineralization. This set of Target Genes is expected to include new drug targets for the treatment of osteoporosis, a common disease for which novel therapeutic options are a health priority.

## Online Methods

### Curating osteoporosis associated outcomes in the UK Biobank study

During the period from 2006 - 2010, half a million British adults were recruited by the UK Biobank study (http://www.ukbiobank.ac.uk/).^42^ Subjects provided biological samples, consented to physical measurements and answered questionnaires relating to general health and lifestyle. Ethical approval was granted by the Northwest Multi-Centre Research Ethics Committee, and informed consent was obtained from all participants prior to participation. Heel bone quality was evaluated in 487,428 subjects by quantitative ultrasound speed of sound (SOS) and broadband ultrasound attenuation (BUA) using a Sahara Clinical Bone Sonometer (Hologic Corporation, Bedford, Massachusetts, USA). Further information regarding the assessment protocols are publicly available on the UK Biobank website. Participants were initially measured at baseline (N = 487,428) and had their left calcaneus (N = 317,815), right calcaneus (N = 4,102) or both calcanei (N = 165,511) measured. A subset of these subjects was followed up at two further time points (N = 20,104 and N = 7,988), during which both heels were measured. A detailed description of the ascertainment procedure is provided in **Figure S1**. Prior to quality control, ultrasound data were available for 488,683 individuals at either baseline and/or follow-up assessment. To reduce the impact of outlying measurements we first identified subjects that had both heels measured and removed those with highly discrepant (i.e. left vs. right) SOS and/or BUA measurements. To achieve this, subjects were stratified by sex and bivariate scatter plots comparing left and right heel measures of SOS and BUA were generated separately. Outliers were identified by manual inspection and removed. The same method was used to identify and remove individuals with highly discordant SOS v BUA measured for each heel. Strict quality control was thereafter applied to male and female subjects separately using the following exclusion thresholds: SOS [Male: (≤ 1,450 and ≥ 1,750 m/s), Female (≤ 1,455 and ≥ 1, 700 m/s)] and BUA [Male: (≤ 27 and ≥ 138 dB/MHz), Female (≤ 22 and ≥ 138 dB/MHz)]. Individuals exceeding the threshold for SOS or BUA or both were removed from the analysis. Estimated bone mineral density [eBMD, (g/cm2)] was derived as a linear combination of SOS and BUA (i.e. eBMD = 0.002592 * (BUA + SOS) − 3.687). Individuals exceeding the following thresholds for eBMD were further excluded: [Male: (≤ 0.18 and ≥ 1.06 g/cm^2^), Female (≤ 0.12 and ≥ 1.025 g/cm^2^)]. A unique list of individuals with a valid measure for the left calcaneus (N = 477,380) and/or right (N = 181,953) were identified separately across the three time points. Individuals with a valid right calcaneus measure were included in the final data set when no left measures were available, giving a preliminary working dataset of N=481,100, (left = 475,724 and right = 5,376) unique individuals. Bivariate scatter plots of eBMD, BUA and SOS were again visually inspected and 723 additional outliers were removed, leaving a total of 480,377 valid QUS measures for SOS, BUA and BMD (264,304 females and 216,073 males). The R script used to curate the raw data is available on request, together with all supporting summary data and plots. Descriptive statistics of the cohort, after quality control, are detailed in **Table S1**.

Fracture cases were identified using two mutually non-exclusive methods: Hospital Episodes Statistics linked through NHS Digital (http://content.digital.nhs.uk/hes) with a hospital-based fracture diagnosis irrespective of mechanism within the primary (N = 392,292) or secondary (N = 320,448) diagnosis field, and questionnaire-based self-reported fracture within the past five years (N = 501,694). We defined a set of International Classification of Diseases codes, 10^th^ revision (ICD10), to separate fracture cases from controls with the Hospital Episodes Statistics data. We excluded fractures of the skull, face, hands and feet, pathological fractures due to malignancy, atypical femoral fractures, periprosthetic and healed fracture codes. A full list of ICD10 codes used can be found in **Table S22**. We did not exclude any self-reported fracture cases by fracture site, since participants were only asked if they sustained a fracture at ankle, leg, hip, spine, write, arm, other or unknown. We identified 20,122 fractures using ICD10 codes and 48,818 using questionnaire-based self-reported data. Descriptive statistics of the cohort, after quality control and ancestry selection, are detailed in **Table S1**.

### Ancestry assignment

Genotype array data were imputed by the UK Biobank using the Haplotype Reference Consortium (HRC) panel^43^. A comprehensive description of the imputation protocol is described elsewhere^44^. A sample of 409,728 White-British individuals was identified centrally by the UK Biobank, using a combination of self-reported ethnicity and genetic information. However, the reliance on self-reported information was deemed too conservative and we chose to redefine a White-British sample (N = 440,414) using genetic information only. We projected the UK Biobank sample onto the first 20 principal components estimated from the 1000 Genomes Phase 3 (1000G) project data^45^ (where ancestry was known) using FastPCA version 2.^46^ Projections used a curated set of 38,551 LD-pruned HapMap 3 Release 3 (HM3)^47^ bi-allelic SNPs that were shared between the 1000G and UK Biobank datasets (i.e. MAF > 1%, minor allele count > 5, genotyping call rate > 95%, Hardy-Weinberg P > 1×10^−6^, and regions of extensive LD removed). Expectation Maximization (EM) clustering (as implemented in R using EMCluster^48^) was used to compute probabilities of cluster membership based on a finite mixture of multivariate Gaussian distributions with unstructured dispersion. Eigenvectors 1, 2 and 5 were used for clustering as they represented the smallest number of eigenvectors that were able to resolve the British 1000G sub-population (GBR) from other ethnicities (**Figure S20**). Twelve predefined clusters were chosen for EM clustering as sensitivity analyses suggested that this number provided a good compromise between model fit (as quantified by log likelihood, Bayesian information criterion, and Akaike information criterion) and computational burden (**Figure S21**). UK Biobank participants (N = 440,414) that clustered together with the 1000G GBR sub-population were termed White-British and used for downstream genetic analyses (**Figure S22**).

### Identification of unrelated samples for LD reference estimation and X chromosome analyses

Genome-wide complex trait analysis (GCTA)^49^ was used to construct a genetic relatedness matrix (GRM) using the White-British sample and a curated set of LD non-pruned HM3 autosomal genome-wide variants (N = 497,687). Unrelated individuals were defined using the genome-wide relatedness measure defined by Yang *et al*.^49^ where the pairwise relatedness between individuals *j* and *k* (*A_jk_*) was estimated by:

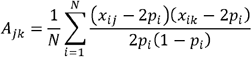

where *x_ij_* is the number of copies of the reference allele for the *i*^th^ SNP of the *j*^th^ and *k*^th^ individuals and *p_i_* is the frequency of the reference allele across the N individuals.

Two samples of unrelated individuals were defined from the White-British UK Biobank population: A sample used for X chromosome association analysis (pairwise relatedness < 0.1, N = 374,559) and a random sample for LD reference estimation (pairwise relatedness < 0.025, N = 50,000).

### Genome-wide association analysis

A maximum of 426,824 White-British individuals (233,185 females and 193,639 males) with genotype and valid QUS measures were analyzed (**Table S1**). For fracture, a maximum of 426,795 White-British individuals, comprising 53,184 fracture cases (60% female) and 373,611 controls (54% female) were analyzed. We note that the sample sizes between the two assessed traits are similar but different, due to not all fracture cases and controls having eBMD measured, and vice-versa. We tested autosomal genetic variants for association with eBMD and fracture, separately, assuming an additive allelic effect, using a linear mixed non-infinitesimal model implemented in the BOLT-LMM v2 software package^50^ to account for population structure and cryptic relatedness. The following covariates were included as fixed effects in all models: age, sex, genotyping array, assessment center and ancestry informative principal components 1 to 20. Autosomal analysis was restricted to up to 13,977,204 high quality HRC imputed variants with a MAF > 0.05%, minor allele count > 5, info score > 0.3, genotype hard call rate > 0.95, and Hardy-Weinberg equilibrium *P* > 1×10^−6^. We also analyzed the association between eBMD and fracture and directly genotyped SNPs on the X chromosome, adjusting for the same covariates, using the Plink2 (October 2017) software package^51^ and a nested sample of unrelated participants (N = 362,926 for eBMD and N = 45,087 cases and 317,775 controls for fracture). As the analyses for the X chromosome data were based upon observed genotypes, we excluded SNPs with evidence of deviation from Hardy-Weinberg Equilibrium (P < 1×10^−6^), MAF < 0.05%, minor allele count ≤ 5, and overall missing rate > 5%, resulting in up to 15,466 X chromosome SNPs for analysis. Heterogeneity in effect size coefficients between sexes was tested in EasyStrata^52^, using Cochran’s test of heterogeneity^53^

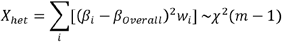

*β_i_ effect size estimates of stratum i*

*SE_i_ standard error of stratum i*

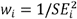

*i = 1..m*

Manhattan plots of our genome-wide association scans were generated using the same software. We have previously estimated the genome-wide significance threshold α = 6.6×10^−9^ for analyzing data from the UK Biobank using the above critera.^4^

### Fracture replication meta-analysis

14 genome-wide significant conditionally independent lead SNPs identified from our fracture analyses were tested for replication in the 23andMe cohort. Genetic associations were tested against the fracture phenotype on a set of unrelated individuals of European ancestry. Analyses were adjusted for age, sex, principal components 1 to 5, and the genotyping platform. There were 367,900 cases and 363,919 controls. Meta-analysis of UK Biobank discovery and 23andMe replication data was performed using METAL.^54^ In order to compare the effect estimates and standard errors of the UK Biobank discovery and 23andMe replication data, we had to transform the UK Biobank discovery effect estimates and standard errors as per the manual specifications in the BOLT-LMM^50^ documentation, specifically:

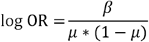

where *μ* = case fraction and standard errors of SNP effect estimates should also be divided by (μ * (1 − μ)).

### Approximate conditional association analysis

To detect multiple independent association signals at each of the genome-wide significant eBMD and fracture loci, we applied approximate conditional and joint genome-wide association analysis using the software package GCTA v1.91.^19^ Variants with high collinearity (multiple regression *R*^2^ > 0.9) were ignored and those situated more than 20 Mbp away were assumed to be independent. A reference sample of 50,000 unrelated White-British individuals randomly selected from the UK Biobank was used to model patterns of linkage disequilibrium (LD) between variants. The reference genotyping dataset consisted of the same variants assessed in our GWAS. Conditionally independent variants reaching genome-wide significance were annotated to the physically closest gene using Bedtools v2.26.0^55^ and the hg19 gene range list (www.cog-genomics.org/plink2).

### Estimation of variance explained by significant variants and SNP heritability

We estimated the proportion of eBMD phenotypic variance tagged by all SNPs on the genotyping array (i.e. the SNP heritability) using BOLT-REML^50^ and Linkage Disequilibrium Score Regression (LDSC)^56^. To calculate the variance explained by independent genome-wide significant SNPs, i.e. all 1,103 genome-wide significant conditionally independent lead SNPs, we summed the variance explained per SNP using the formula: 2p(1 - p)β^2^, where p is the effect allele frequency and β is the effect of the allele on a standardized phenotype (mean = 0, variance = 1).^57–59^

### Estimating genomic inflation with LD Score Regression (LDSC)

To estimate the amount of genomic inflation present in the data that was due to residual population stratification, cryptic relatedness, and other latent sources of bias, we used stratified LDSC^60^ in conjunction with partitioned LD scores that were calculated for high quality HM3 SNPs derived from a sample of unrelated 1000G EUR individuals.

Fine-Mapping SNPs

Fine-mapped SNPs were defined as those being conditionally independent, as identified by GCTA-COJO or exceeding our threshold for posterior probability of causality, as defined by FINEMAP. Here we describe the generation of this set of fine-mapped SNPs.

First, SNPs were defined as being conditionally independent using GCTA-COJO.^19,20^ We next calculated the posterior probability of causality. To do so, we defined each conditionally-independent lead SNP as a signal around which, we would undertake posterior probability testing. We used all imputed SNPs within 500 kbp of a conditionally independent lead SNP and treated each signal independently. We used FINEMAP^20^, which approximates, per input region, genotype-phenotype data with correlation matrices and summary statistics, and then implements a shotgun stochastic search algorithm to test causal configurations of SNPs rapidly and identify the most likely number of causal SNPs per signal in a Bayesian framework. We generated correlation matrices for each fine-mapped region from a subset of randomly selected 50,000 White-British UK Biobank participants with the LDSTORE software^61^. FINEMAP was run with default parameters except for the number of maximum causal configurations tested, which we set to 10.^20^ For the causal configuration with the highest posterior probability, each SNP was assigned a log_10_ Bayes factor as a measure of its posterior probability for being in the causal configuration. For example, if a tested region had a causal configuration of six SNPs with the highest posterior probability, all tested SNPs were assigned a Bayes factor for their marginal posterior probabilities of being in that causal configuration. Based on this information we constructed our sets of fine-mapped SNPs, including only the SNPs with the highest posterior probabilities. After testing each signal at a locus, the set of fine-mapped SNPs were collapsed into the same locus, due to the high amount of redundancy between credible sets for each signal, given that the approximation of genotype-phenotype data with correlation matrices and summary statistics implemented by FINEMAP is identical to GCTA-COJO.^19,20^ We used a log_10_ Bayes factor > 3 threshold to only consider SNPs with the strongest posterior probabilities for causality, and those SNPs that were identified as genome-wide significant conditionally independent lead SNPs, as being fine-mapped SNPs.

### RNA sequencing for mouse osteocytes

We performed an analysis of whole transcriptome sequencing data of three distinct bone types from the mouse skeleton to measure osteocyte expression^4^. The three sites were the tibia, femur and humerus, and in each, the bone marrow was removed (N = 8 per site). The distribution of normalized gene expression for each sample was used to calculate a threshold of gene expression^62^, with genes above this threshold for 8 out of 8 replicates in any bone type deemed to be expressed. Osteocyte enriched genes were determined by comparing the transcriptomes of matched bone sample controls, one with the marrow removed and the other with the marrow left intact (N = 5 per site). Genes significantly enriched in osteocytes and expressed in all bone types were defined as osteocyte transcriptome signature genes.

### Mapping accessible chromatin

ATAC-seq libraries were generated by the McGill University and Genome Quebec Innovation Centre on 100,000 SaOS-2 cells, using a modified protocol to that previously described^63^. The modifications included: reducing the transposase reaction volume from 50 μl to 25 μl, increasing the transposase concentration from 1x to 40x, and using 12 cycles of PCR to enrich each library. Libraries were quantified by Q-PCR, Picogreen and LabChip, then were sequenced on the Illumina HiSeq 2500 to 125 bp in pair-ended mode, using the Nextera sequencing primers. DNase-seq data from primary osteoblast samples^21^ were obtained from http://encodeproject.org under accessions ENCLB776DWN and ENCLB906BCL.

Reads were processed using a uniform pipeline to produce both ATAC-seq and DNase-seq peaks. Illumina adapters were trimmed using Trimmomatic v. 0.36^64^. Reads were aligned to the hg38 human reference using BWA v.0.7.15 65. Peak calling was performed using hotspot2 (https://github.com/Altius/hotspot2) with a cutoff of 1% FDR and converted to hg19 reference coordinates using UCSC liftOver.

### RNA sequencing for human osteoblast cell lines

RNA library preparations were carried out on 500□ng of RNA from SaOS-2, U2OS, MG63 and HOS cells with RNA integrity number (RIN) > 7 using the Illumina TruSeq Stranded Total RNA Sample preparation kit, according to manufacturer’s protocol. Final libraries were analyzed on a Bioanalyzer and sequenced on the Illumina HiSeq4000 (pair-ended 100□bp sequences). Raw reads were trimmed for quality (phred33 ≥ 30) and length (n ≥ 32), and Illumina adapters were clipped off using Trimmomatic v. 0.35^64^. Filtered reads were aligned to the GRCh37 human reference using STAR v. 2.5.1b^65^. Raw read counts of genes were obtained using HTseq-count v.0.6.1^66^.

### RNA sequencing for murine calvarial osteoblasts

We used whole transcriptome sequencing on mouse osteoblasts post-differentiation to obtain expression profiles of the maturing osteoblast^4^. We obtained pre-osteoblast-like cells from the neonatal calvaria of C57BL/6J mice carrying a Cyan Fluorescent Protein (*CFP*) transgene under the control of the *Col* 3.6 kbp promoter^67^. Specifically, we removed cells not expressing CFP by FACS sorting after culturing for four days in growth media. The remaining cell set was considered enriched for pre-osteoblast cells and was re-plated and subjected to an osteoblast differential cocktail, with RNA being collected every two days from days two to 18 post-differentiation. We used whole transcriptome sequencing with three technical replicates per sample and calculated a normalized expression level per gene.

### High-throughput chromosome conformation capture

High-throughput chromosome conformation capture (Hi-C) was performed on primary human osteoblasts and osteocytes from human bone biopsies of non-fracture subjects. Hi-C libraries were prepared as described previously.^68^ Instead of using HindIII restriction enzyme, we used DpnII^69^ which increased coverage and insensitivity of CpG methylation^70^. The Hi-C libraries were sequenced on Illumina HiSeq4000 instruments to 2 billion pair-end reads. Biological replicates were independently generated and sequenced. HiC-Pro was used to process the HiC-Pro pipeline^71^ beginning with aligning each read end to hg38 reference genomes. The Chimeric read ends were filtered to keep only 5’ alignments with MAPQ > 10, and then read-ends were paired and de-duplicated. Contact matrices were constructed, and significant interactions were estimated with Homer,^72^ GOTHiC^73^ and Juicer.^74^ We defined significant interactions as P < 10^−15^ (comparing observed interactions to estimated expected interactions and taking into account DNA fragment size, GC content, and other genomic features). Only interaction pairs that were significant (P < 10^−15^) from all three tools were considered significant. The resolution of Hi-C interactions was from 1.5 to 2 kbp with average 1.8 kbp. ATAC-seq experiments were also performed in primary osteoblasts and osteocytes that were used for HI-C experiments. We only considered and reported chromatin interactions that mapped to open chromatin

### Target Gene identification

We identified Target Genes for the autosomal fine-mapped sets by annotating fine-mapped sets of SNPs to the closest protein-coding gene, making additional note if the SNP mapped directly to the gene’s introns or exons, or was coding. We identified Target Genes on the X chromosome by the closest gene to a conditionally independent lead SNP, as we did not calculate log_10_ Bayes factors for SNPs on the X chromosome. Additionally, we annotated Target Genes that may be functional in bone cells by marking which fine-mapped SNPs mapped to open chromatin in human bone cells, identified by SaOS-2 ATAC-seq peaks, and we mapped chromosomal positions of fine-mapped SNPs to significant Hi-C interactions of primary osteoblast and osteocytes. When the interaction chromatin mapped to multiple isoforms of protein coding genes, we selected the one with the most significant interaction (usually with highest interaction counts). When the interaction chromatin mapped to multiple bins, we selected the one(s) with looping domains. We further annotated Target Genes using the osteocyte signature gene set where genes within this set are enriched for osteocyte activity.^4^

### Target Gene enrichment analyses

We performed a series of enrichment analyses by calculating the odds of Target Genes being either positive control genes or osteocyte signature genes. We identified a set of 57 proteins whose perturbation through pharmacotherapy,^2^ or Mendelian disease leads to changes in bone density, monogenic disorders presenting with abnormal skeletal mineralization or low bone mass, osteolysis and/or skeletal fragility and osteogenesis imperfecta and abnormal skeletal mineralization (**Table S12**).^22^ For all protein-coding genes in the genome, which were identified using refGene (N = 19,455), we annotated whether they were found to be Target Genes and/or positive control genes. These annotations allowed us to construct contingency tables and calculate an odds ratio for enrichment of Target Genes amongst positive control genes. We used multiple genomic features to test which methods of identifying Target Genes enriched for positive control genes. To do so, we tested if positive control genes were enriched amongst targeted genes identified by four different methods: 1) Genes that were most proximal to the fine-mapped set SNPs; 2) Genes that contained fine-mapped SNPs overlapping their gene bodies; 3) Genes containing fine-mapped SNPs that are coding variants; 4) Genes identified to be in 3D contact with fine-mapped sets in human osteoblasts or osteocytes through Hi-C experiments; 5) The closest gene to fine-mapped SNPs, which also mapped to ATAC-seq peaks in human osteoblast SaOS-2 cell lines; and 6) Those genes within 100 kbp of fine-mapped SNPs (Figures 2 and 4). We then repeated this analysis using the osteocyte signature gene set (N = 1,240) instead of the positive control set, to calculate the odds of Target Genes being active in the osteocyte.

### Target Gene pathway analysis

We used the Functional Mapping and Annotation of GWAS tool (FUMA)^23^ to annotate our lists of Target Genes for their most enriched biological pathways with data from the WikiPathways^24^ database. WikiPathways is an openly curated database for biological pathways and provides information on the roles of specific genes or proteins in their respective pathways. FUMA uses WikiPathways data to compare a list of given genes against a background gene set (e.g. all protein coding genes) with hypergeometric testing. The output is then a list of enriched biological pathways based on the input gene lists. We have presented these data graphically in the **Figure S7**.

### CRISPR/Cas9 Methods

SaOS-2 cells were obtained from ATCC (#ATCC HTB-85) and cultured in McCoy5A medium (ATCC) supplemented with 15% of FBS (Wisent inc) and 1% of penicillin and streptomycin (Wisent Inc.) according to the manufacturer. Three different guide RNAs (gRNA) targeting the second exon of *DAAM2* were cloned in the PX458 plasmid (pSpCas9(BB)-2A-GFP; Addgene #48138). The gRNA sequences were: gRNA 1-CAGAGGGTGGTTGTCCCGG; gRNA 2-CAGCCCCATCCCGAACGCAG; and gRNA 3-TGTCCCGGAGGTTGATTTCG. We observed the cutting frequency determination (CFD) scores^75^ for each gRNA was < 0.1, therefore we did not consider off-target effects to merit testing^76^. The construct plasmids were purified using the QIAGEN filter midi prep kit (QIAGEN #12243) according to manufacturer instructions. SaOS-2 cells were cultured to 80% confluence in a 100-mm^2^ petri dish. Cells were then transfected with one of the three different plasmids generated, or with the intact plasmid as a control, using TransIT LT1 transfection reagent (Mirus #MIR2304) with a reagent-to-DNA ratio of 3:1. 48 hours post-transfection, GFP positive cells were sorted by FACS in a single cell model. The remaining colonies were expanded and then assessed for the presence of DAAM2 protein using immunofluorescence technique (Anti-DAAM2 antibody, Sigma-Aldrich #HPA051300). PCR primers were designed against regions of *DAAM2* flanking the three gRNA target sequences (forward: 5′-tcctcttgtccagATCACAATG-3′ and reverse: 5′-ccaagaggagttttgagagatgga-3′) to generate an amplicon of 355 bp. PCR products of the identified clones were sequenced using MiSeq (Genome Quebec).

To generate DAAM2 Western blots (**Figure S23**), total protein was extracted from SaOS-2 cells using a RIPA buffer. Denatured proteins (20 μg) were separated by 10% sodium dodecylsulfate (SDS) polyacrylamide gel electrophoresis followed by transfer to nitrocellulose membranes. The membranes were blocked in 5% skim milk for one hour at room temperature followed by incubation with an anti-DAAM2 antibody (Abcam #ab169527) at 1/1,000 overnight at 4°C and the secondary antibody goat anti-rabbit IgG at 1/10,000 for one hour at room temperature (Abcam #ab205718). The band densities were quantified by densitometry using Image Lab 5.1 software (Bio-Rad). Protein levels were expressed as a ratio of protein-specific band density and that of total protein stained using MemCode Staining Solution (Thermofisher #24580). **Figure S23** shows that DAAM2 protein expression was reduced to 17.5% and 33.5% in the gRNA1 and gRNA2 edited clones, respectively.

To induce mineralization (Figure 5), cells were then cultured to 90% confluence in a 6-well plate and then treated, or left untreated for a control, with osteogenic factors (Ascorbic acid 50 μg/ml and β-Gycerophosphate 10 mM). Fresh media containing osteogenic factors was added every 2-3 days over 13 days. At day 14, mineralization was quantified using the osteogenesis assay kit according to manufacturer instructions (Millipore #ECM815). The Alizarin red concentration (μM) was normalized with the protein content assessed in the media in each culture (Pierce BCA Protein assay kit; Thermo Fisher #23227).

### Rapid throughput murine knockout program

The Origins of Bone and Cartilage Disease (OBCD) program (www.boneandcartilage.com) is undertaking rapid-throughput structural and functional skeletal phenotype analyses of all unselected knockout mice generated at the Wellcome Trust Sanger Institute as part of the International Knockout Mouse and International Mouse Phenotyping Consortia (IKMC and IMPC). Anonymized samples from 16-week-old female wild-type and mutant mice (N = 2 to 6 per mutant genotype) were stored in 70% ethanol and assigned to batches for rapid throughput analysis. Mice were fed either a Breeder’s Chow (Mouse Breeder Diet 5021, 21% kcal as fat, Labdiet, London, UK) or a Western diet (Western RD, 829100, 42% kcal as fat, Special Diet Services, Witham, UK) from 4 weeks of age. The relative bone mineral content and length of the femur and caudal vertebrae are determined by digital X-ray microradiography (Faxitron MX20, 10μm pixel resolution)^77–79^. Micro-CT (Scanco uCT50, 70kV, 200μA, 0.5mm aluminium filter) is used to determine trabecular parameters (bone volume BV/TV, trabecular number Tb.N, thickness Tb.Th, spacing Tb.Sp) at a 5μm voxel resolution in a 1mm region beginning 100μm proximal to the distal femoral growth plate and cortical bone parameters (thickness Ct.Th, BMD, medullary diameter) at a 10μm voxel resolution in a 1.5mm region centered in the mid-shaft region 56% along the length of the femur distal to the femoral head.^77,80,81^ Biomechanical variables of bone strength and toughness (yield load, maximum load, fracture load, % energy dissipated prior to fracture) are derived from destructive 3-point bend testing of the femur and compression testing of caudal vertebra 6 and 7 (Instron 5543 load frame, 100N and 500N load cells).^77,79^ Overall, 19 skeletal parameters were reported for each individual mouse studied and compared to reference data obtained from 320 16-week-old wild-type C57BL/6 female mice. Outlier phenotypes were defined by parameters > 2 standard deviations away from the reference mean determined from the 320 age, sex and genetically identical C57BL/6N wild-type controls. Enrichment of outlier skeletal parameters in mice with deletion of eBMD Target Genes was determined by comparison with the frequency of outlier parameters in 526 unselected knockout lines using Fisher’s Exact Test (**Table S18**, Prism, GraphPad Software, La Jolla, USA). The 526 unselected knockout lines were generated by the WTSI and phenotyped by the OBCD program; these lines included 56 Target Genes. Five Target Genes had previously been phenotyped in an OBCD pilot study^77^ and knockout lines for an additional 65 Target Genes, that had already been generated by WTSI, were prioritized for rapid-throughput skeletal phenotyping. In total, our analyses included 596 knockout lines.

Additional skeletal samples from 16-week-old WT (n=5 female, n=5 male), *Daam2^+/tm1a^* (n=7 female, n=5 male) and *Daam2^tm1a/tm1a^* (n=7 female, n=5 male) mice were analyzed as described above. Supplementary cortical bone parameters (total cross-sectional area Tt.Ar, cortical bone area Ct.Ar, medullary area M.Ar, periosteal perimeter Ps.Pm, endocortical perimeter Ec.Pm, cortical porosity Ct.Po, polar moment of inertia (*J*) and maximum and minimum moments of inertia (/max and /min)) were determined by micro-CT (at 10μm voxel resolution, except for Ct.Po which was determined at 1μm voxel resolution using the Scanco uCT50 at 70kV, 57μA, 0.5mm aluminium filter). Correlation between bone mineral content and biomechanical parameters was determined by linear regression analysis using 320 16-week-old WT femur and vertebra samples from C57BL/6 female mice. Bone composition and structure was investigated in *Daam2^tm1a/tm1a^* mice by comparing observed biomechanical parameters with values predicted by linear regression analysis of femoral and vertebral BMC and biomechanical parameters obtained from 320 WT age and sex matched controls.

### *Daam2* knockout mice

Mouse studies undertaken at the Garvan Institute of Medical Research (Darlinghurst, NSW, Australia) were approved by the Garvan Institute / St Vincent’s Hospital Animal Ethics Committee in accordance with New South Wales (Australia) State Government legislation. *Daam2^tm1a(KOMP)Wtsi^* mice (designated *Daam2^tm1a/tm1a^*) were obtained from the Wellcome Trust/Sanger Institute (Cambridge, UK) where the mice were generated as part of the International Mouse Phenotyping Consortium (http://www.sanger.ac.uk/mouseportal), using ES cells produced by the Knockout Mouse Project (https://www.komp.org/geneinfo.php?Symbol=Daam2). The *Daam2* gene in these mice was disrupted by a cassette containing an insertion with an additional splice acceptor site between exons 5 and 6 (http://www.mousephenotype.org/data/alleles/MGI:1923691/tm1a%28KOMP%29Wtsi?). The success of this strategy was confirmed with an 80% knockdown of *Daam2* in *Daam2^tm1atm1a^* and 50% knockdown in *Daam2^+/tm1a^*. Age and sex matched 16-week old mice were used for detailed skeletal phenotyping, as described above.

### In vitro assays of osteoclast formation

Osteoclasts were generated from primary BMCs flushed from mouse long bones of 8-10 week old WT, *Daam2^+tm1a^* and *Daam2^tm1a/tm1a^* mice, resuspended in MEM/FBS then added (10^5^ cells/well) to 6mm diameter culture wells. These were stimulated with 10, 20, 50 and 100 ng/ml RANKL, plus 50 ng/mL M-CSF. Medium and cytokines were replaced at day 3, and on day 6 cultures were fixed with 4% paraformaldehyde and histochemically stained for TRAP using as previously described.^82^ TRAP positive multinucleated cells (MNCs) containing 3 or more nuclei were counted as osteoclasts and quantified under inverted light microscopy.

### In vitro osteoblast mineralization

Plastic-adherent bone marrow stromal cells (BMSCs) were isolated from 8-10 week old WT, *Daam2^+tm1a^* and *Daam2^tm1a/tm1a^* mice as described previously. ^83^ Briefly, marrow cells were flushed from mouse long bones and plated in MEM containing 20% FBS in 25cm^2^ tissue culture flask. Non-adherent cells were removed by medium changes 3 and 5 days later. After 7 days in culture, cells were trypsinized, scraped and re-plated at 3 x 10^4^ cells/cm2 in 24-well plates in MEM with 10% FBS containing osteoblast differentiating factors (50 μg/ml ascorbic acid, 2.5nM dexamethasone and 10 mM β-glycerolphosphate; Sigma-Aldrich), which was added and changed every 3 days for 21 days. Cells were washed with PBS and fixed with 4% paraformaldehyde for 15 mins then ethanol (80%) for 30 mins, rinsed and stained with 0.5% Alizarin Red (Sigma Aldrich) in water for 30 mins, washed, dried and images of the plates taken with a flat-bed scanner (model v800, Epson, North Ryde, NSW Australia). Alizarin red was then eluted with 10% cetyl pyridinium chloride (CTP; Sigma-Aldrich) in PBS overnight and quantified by measuring 562 nm absorbance (Clariostar plate reader, BMG Labtech, Offenburg, Germany) relative to standard alizarin red solutions.

### Detection of serum markers of bone resorption and formation

Serum levels of bone resorption marker tartrate-resistant acid phosphatase (TRAP) and bone formation marker procollagen type 1 N-terminal propeptide (P1NP) were measured using a Rat/Mouse TRAP enzyme immunoassay kit and a Rat/Mouse P1NP enzyme immunoassay kit (Immunodiagnostic Systems, Gaithersburg, MD, USA) respectively.

### Fourier-Transform Infrared Spectroscopy

The humeri from *Daam2* WT, *Daam2^+/tm1a^* and *Daam2^tm1a/tm1a^* male and female mice were collected at 16 weeks of age. 21 male samples (11 WT, 4 *Daam2^+/tm1a^* and 6 *Daam2^tm1a/tm1a^)* and 19 female samples (8 WT, 5 *Daam2^+/tm1a^* and 6 *Daam2^tm1a/tm1a^*) were examined. The bones were frozen immediately and were kept at a stable temperature until analysis. All bones were processed at the same time and all analyzed on the same day to reduce batch effects. The humeri were thawed, stripped of soft tissue with epiphyses removed and the marrow cavity was flushed. Specimens were then refrozen in liquid nitrogen and pulverized at −80°C using a SPEX Sample Prep 6870 Freezer/Mill. Each sample was subjected to three rounds of pulverization at 15 cycles per second for one minute with a two-minute cool-down between each round. The samples were lyophilized under vacuum at −51°C overnight to ensure they were completely dehydrated. Anhydrous potassium bromide (KBr) was then added until the final concentration of bone in the samples was between 2.50-2.56% by mass. KBr pellets were formed by compressing 20 mg of mixed KBr and bone samples in a 7 mm die under 4 tons of force. The formed pellets were loaded into a Nicolet iS50 FT-IR spectrophotometer (Thermo Fisher Scientific). The collection chamber was continuously purged with dry nitrogen gas to minimize noise from moisture and carbon dioxide. Background noise was collected on KBr-only pellets and subtracted at the beginning of each cohort or after 30min of continuous measurements (whichever occurred first). For each sample, 128 scans between 400-2200 cm^−1^ (wave numbers) were collected at a resolution of 4.0cm^−1^ using Happ-Genzel apodization. The wave number data was curve fit to absorbance, with baselining and spectral analyses performed using custom algorithms and scripts written in the R programming language (R version 3.4.2). The scripts were built on top of the ChemoSpec (version 4.2.8) and MESS (version 0.3-2) packages. Local minima were used as limits of integration to calculate areas under the curve for the carbonate, phosphate and amide I peaks; the mineral to matrix ratio, carbonate to phosphate ratio were then calculated using these areas in the appropriate ratios. Collagen maturity and crystallinity were calculated from the spectral data using absorbance values at literature-reported and validated wavenumbers.^84^ Between two and four technical replicates were run for each sample, based on the amount of material available. Two samples (both from WT males) were removed from all subsequent statistical analyses as the signal to noise ratio was excessive for the spectral data for all technical replicates, thus precluding obtaining meaningful information from those samples. Values for technical replicates where averaged for each specimen. Differences between genotypes were determined by ANOVA, followed by a Tukey’s post hoc test. Data from male and female mice were analyzed separately.

### URLs

URLs to download the genome-wide association summary statistics for eBMD and fracture, as well as RNA-seq and ATAC-seq data generated for human osteoblast cell lines, will be made available after peer-reviewed publication.

---

**Figure S1. Flow diagram illustrating calcaneal quantitative ultrasound (QUS) data collection by the UK Biobank**. QUS data were collected at three time points: Baseline (2007 - 2010), Follow-up 1 (2012 - 2013) and Follow-up 2 (2014 - 2016). At baseline, QUS was performed using two protocols (denoted protocol 1 and 2). Protocol 1 was implemented from 2007 to mid-2009 and involved measuring the left calcaneus. Only in cases where the left was missing or deemed unsuitable was the right calcaneus measured. Protocol 2 was introduced from mid-2009, (replacing protocol 1) and differed only in that it involved measuring both the left and right calcanei. Protocol 2 was further used for both follow up assessments. For all three time points, calcaneal QUS was performed with the Sahara Clinical Bone Sonometer [Hologic Corporation (Bedford, Massachusetts, USA)]. Vox software was used to automatically collect data from the sonometer (denoted direct input). In cases where direct input failed, QUS outcomes were manually keyed into Vox by the attending healthcare technician or nurse (i.e. manual input). The number of individuals with non-missing measures for speed of sound (SOS) and broadband ultrasound attenuation (BUA) recorded at each assessment period are indicated in light grey. Further details on these methods are publicly available on the UK Biobank website (UK Biobank document #100248 https://biobank.ctsu.ox.ac.uk/crystal/docs/Ultrasoundbonedensitometry.pdf). To reduce the impact of outlying measurements, Individuals with highly discordant left vs. right calcaneal measures were excluded from the analysis. Furthermore, quality control was applied to male and female subjects separately using the following exclusion thresholds: SOS [Male: (≤ 1,450 and ≥ 1,750 m/s), Female (≤ 1,455 and ≥ 1,700 m/s)] and BUA [Male: (≤ 27 and ≥ 138 dB/MHz), Female (≤ 22 and ≥ 138 dB/MHz)]. Individuals exceeding the threshold for SOS or BUA or both were removed from the analysis. Estimated bone mineral density [eBMD, (g/cm2)] was derived as a linear combination of SOS and BUA (i.e. eBMD = 0.002592 * (BUA + SOS) − 3.687). Individuals exceeding the following thresholds for eBMD were further excluded: [Male: (≤ 0.18 and ≥ 1.06 g/cm2), Female (≤ 0.12 and ≥ 1.025 g/cm2)]. The number of individuals with non-missing measures for SOS, BUA and eBMD after QC are indicated in black. A unique list of individuals with a valid measure for the left calcaneus (N=477,380) and/or right (N=181,953) were identified separately across all three time points. Individuals with a valid right calcaneus measure were included in the final data set when no left measures were available, giving a preliminary working dataset of N=481,100, (left=475,724 and right=5,376) unique individuals. Bivariate scatter plots of eBMD, BUA and SOS were visually inspected and 723 additional outliers were removed, leaving a total of 480,377 valid QUS measures for SOS, BUA and BMD (264,304 females and 216,073 males).

**Figure S2. QQ plot of GWAS for eBMD**. Results are from the entire genome and not conditionally independent SNPs.

**Figure S3. Relationship between absolute effect size (y-axis) and minor allele frequency (x-axis) for 1,103 conditionally independent SNPs**. Red dots represent SNPs at previously reported BMD loci. Blue dots represent SNPs at novel loci. The named gene is that closest to the SNP that has the smallest P-value of all conditionally independent SNPs present in the same locus. We emphasize that proximity is not necessarily indicative of causality.

**Figure S4. Effect size in standard deviations for eBMD (y-axis) from the current UK Biobank Study plotted against effect size in standard deviations from the previous GEFOS studies for BMD at the (A) femoral neck, (B) lumbar spine, (C) forearm, (D) total-body, (E) heel and (F) fracture as per the full UK Biobank cohort (x-axis)**. Only conditionally independent variants that reach genome-wide significance (P < 6.6 x10^−9^) for eBMD in the UK Biobank are plotted. Minus log_10_ P-value for the fracture analysis in UK Biobank is represented by the shading of the data points (black for robust evidence of association with fracture and white for poor evidence of association). The blue dashed trend line shows a moderate to strong correlation between estimated effect sizes at the heel and femoral neck [r=0.53 95%-CI (0.49,0.57)], lumbar spine [0.59 (0.55,0.63)], forearm [0.46 (0.41, 0.50)], total-body [0.70 (0.67,0.73)], interim heel [0.93 (0.92,0.94)] and fracture [−0.77 (−0.79, −0.74)]. SNPs that reach genome-wide significance for fracture look-up (P < 6.6 x10^−9^) are labelled in black.

**Figure S5. Manhattan plot of genome-wide association results for fracture in the UK Biobank**. Manhattan plot showing genome-wide association results for fracture in the UK Biobank. The dashed red line denotes the threshold for declaring genome-wide significance (6.6×10^−9^). In total, 14 conditionally independent SNPs at 13 loci passed the criteria for genome-wide significance in the discovery analysis. Blue dots represent a locus identified from the eBMD GWAS that was novel in this analysis. Red dots represent a locus associated with eBMD which was known from previous studies. Previously reported BMD loci failing to reach genome-wide significance in our study are shown in black.

**Figure S6. Analysis of sex heterogeneity for eBMD**. The top-most figure is a Miami plot of genome-wide association results for females (top panel) and males (bottom panel). The bottom graph is a Manhattan plot for the test for sex heterogeneity in eBMD regression coefficients between males and females. Previously reported loci that reached genome-wide significance (P < 6.6 x10^−9^) are displayed in red, and previously reported loci failing to reach genome-wide significance in our study are shown in black. Loci containing *ABO, FKBP4, LOC105370177* and *FAM9B* had stronger effects on eBMD in males, whereas *MCM8* had a larger effect in females. Loci demonstrating significant heterogeneity (P < 6.6 x10^−9^) but were not robustly associated at genome-wide significance with eBMD in the males and/or females are in green (i.e. *MCCD1* and *ZNF398)*.

**Figure S7**. WikiPathways pathway analysis results from FUMA for (**A**) genes closest to a fine-mapped SNP, (**B**) genes with fine-mapped SNPs mapping to its gene body, (**C**) genes with coding fine-mapped SNPs, (**D**) genes mapped closest to a fine-mapped SNP which resided in an SaOS-2 ATAC-seq peak, and genes identified by fine-mapped SNP was present in a (**E**) Hi-C osteoblast or (**F**) osteocyte promoter interaction peak. Well known pathways for bone biology were highlighted by FUMA, such as Wnt signalling, endochondral ossification, osteoclast and osteoblast signalling.

**Figure S8. Expression of *DAAM2* in osteoblast cell lines from RNA Sequencing experiments and open chromatin profiles from ATAC-seq experiments**. Blue shows forward strand expression, while red shows reverse strand expression. Dark purple shows ATAC-seq open chromatin peaks. RNA of *DAAM2* is present in all cell lines, but particularly, SaOS-2, HOS and U-2_OS cell lines.

**Figure S9: No unspecific labeling of the secondary antibody in the SaOS-2 osteoblast cell line**. Representative immunofluorescence of SaOS-2 cell lines stained with goat anti-rabbit IgG Alexa Fluor 488 secondary antibody (Abcam, ab150077; 1/1000), counterstained with DAPI (blue) and observed by confocal microscopy.

**Figure S10. DAAM2 is localized to the nucleus of SaOS-2 osteoblast cell lines**. Representative immunofluorescence of SaOS-2 cell lines stained with anti-DAAM2 antibody (Sigma Aldrich, HPA051300; 1/200) followed by goat anti-rabbit IgG Alexa Fluor 488 secondary antibody (Abcam, ab150077; 1/1000), counterstained with DAPI (blue) and observed by confocal microscopy.

**Figure S11. Additional skeletal phenotyping of *Daam2* knockdown at postnatal day 112. A) Bone mineral content and length**. X-ray microradiography images (Faxitron MX20) showing femur and caudal vertebrae from female (left) and male (right) wild-type (WT; female n=5, male=5), heterozygous *(Daam2^+/tm1a^* female n=7, male n=5) and homozygous (*Daam2^tm1a/tm1a^*; female n=7, male n= 9) knockout mice. Gray scale images of femur and caudal vertebrae are pseudocoloured according to a 16-colour palette in which low mineral content is green and high mineral content is pink. Relative frequency plots showing bone mineral content in femur and caudal vertebrae from WT, *Daam2^+/tm1a^* and *Daam2^tm1a/tm1a^* mice; Kolmogorov–Smirnov test, *P<0.05. Graphs demonstrate femur and caudal vertebra length in WT, *Daam2^+/tm1a^* and *Daam2^tm1a/tm1a^* mice. Data are shown as mean ± SEM; ANOVA and Tukey’s post hoc test; *P<0.05; **P<0.01. **B) Trabecular bone parameters**. Micro-CT images (Scanco MicroCT-50) showing proximal femur trabecular bone from WT, *Daam2^+/tm1a^, Daam2^tm1a/tm1a^* mice. Graphs showing trabecular bone volume/tissue volume (BV/TV), trabecular number (Tb.N), trabecular thickness (Tb.Th) and trabecular spacing (Tb.Sp). Data are shown as mean ± SEM. **C) Cortical bone parameters**. Micro-CT images of mid-diaphysis cortical bone from WT, *Daam2^+/tm1a^, Daam2^tm1a/tm1a^* mice. Graphs showing total cross-sectional area inside the periosteal envelope (Tt.Ar), cortical bone area (Ct.Ar), cortical area fraction (Ct.Ar/Tt.Ar), medullary (marrow cavity) area (M.Ar), periosteal perimeter (Ps.Pm), endocortical perimeter (Ec.Pm), cortical thickness (Ct.Th), cortical bone mineral density (BMD), cortical porosity (Ct.Po), polar moment of inertia (J), maximum moment of inertia (/max) and minimum moment of inertia (/min). Data are shown as mean ± SEM.

**Figure S12: Bone resorption and formation are not affected by Daam2 knockdown. A**) No difference in the number of bone marrow-derived TRAP+ multinucleated cells was observed between WT and *Daam2^tm1a/tm1a^* male mice (Scale bar = 100 μM; n = 4; mean ± SEM). B) No difference was observed in the mineralization of bone marrow stromal cells between WT and *Daam2^tm1a/tm1a^* mice. No difference in bone resorption marker TRAP (C) and bone formation marker P1NP (D) was detected in the sera of WT and *Daam2^tm1a/tm1a^* mice. Data in (C) and (D) are shown as mean ± SEM; Females are on left and males on right.

**Figure S13. Bone composition of *Daam2* knockdown and wildtype mice**. Bone composition was measured in humeri from 16 week old male and female mice by Fourier Transformed Infrared Spectroscopy (FTIR). **A**) Mineral to matrix ratio was determined as the ratio of the integrated areas of the phosphate peak/amide I peak. **B**) Carbonate substitution was defined as the ratio of the integrated areas of the carbonate/phosphate peaks. **C**) Collagen maturity or collagen crosslinking was calculated as the ratio of the peak spectral intensities at 1660 and 1690 cm^−1^ respectively. **D**) Crystallinity or crystal maturity was calculated as the ratio of the peak spectral intensities at 1030 and 1020 cm^−1^ respectively.

**Figure S14**. Increased bone mass, stiffness and strength in adult *Chromobox 1* heterozygous deficient mice (*Cbx1^+/−^*) (a) X-ray microradiography images of femur and caudal vertebrae from female wild-type (WT) and *Cbx1^+/−^* mice at postnatal day 112 (P112). Graphs show reference ranges derived from 320 WT mice, mean (solid line), 1.0SD (dotted lines) and 2.0SD (grey box). Parameters from individual *Cbx1^+/−^* mice are shown as red dots and mean values as a black line (n=2 animals). (b) Micro-CT images of proximal femur trabecular bone (left) and mid-diaphysis cortical bone (right). Graphs showing trabecular bone volume/tissue volume (BV/TV), trabecular number (Tb.N), trabecular thickness (Tb.Th), trabecular spacing (Tb.Sp), cortical thickness (Ct.Th), internal cortical diameter and cortical bone mineral density (BMD). (c) Graphs showing yield load, maximum load, fracture load, stiffness and energy dissipated prior to fracture derived from 3-point bend testing of femurs. (d) Graphs showing yield load, maximum load and stiffness derived from compression testing of vertebra. Scale bars: a, 1mm and b, 0.5mm.

**Figure S15**. Increased bone mass and strength in adult *WW Domain Containing Adaptor with Coiled-Coil* heterozygous deficient mice (*Wac^+/−^*) (a) X-ray microradiography images of femur and caudal vertebrae from female wild-type (WT) and *Wac^+/−^* mice at postnatal day 112 (P112). Graphs show reference ranges derived from 320 WT mice, mean (solid line), 1.0SD (dotted lines) and 2.0SD (grey box). Parameters from individual *Wac^+/−^* mice are shown as red dots and mean values as a black line (n=2 animals). (b) Micro-CT images of proximal femur trabecular bone (left) and mid-diaphysis cortical bone (right). Graphs showing trabecular bone volume/tissue volume (BV/TV), trabecular number (Tb.N), trabecular thickness (Tb.Th), trabecular spacing (Tb.Sp), cortical thickness (Ct.Th), internal cortical diameter and cortical bone mineral density (BMD). (c) Graphs showing yield load, maximum load, fracture load, stiffness and energy dissipated prior to fracture derived from 3-point bend testing of femurs. (d) Graphs showing yield load, maximum load and stiffness derived from compression testing of vertebra. Scale bars: a, 1mm and b, 0.5mm.

**Figure S16**. Increased bone mineral content and strength in adult *DNA Replication and Sister Chromatid Cohesion 1* heterozygous deficient mice (*Dscc1^+/−^*) (**a**) X-ray microradiography images of femur and caudal vertebrae from female wild-type (WT) and *Dscc1^+/−^* mice at postnatal day 112 (P112). Graphs show reference ranges derived from 320 WT mice, mean (solid line), 1.0SD (dotted lines) and 2.0SD (grey box). Parameters from individual *Dscc1^+/−^* mice are shown as red dots and mean values as a black line (n=2 animals). (**b**) Micro-CT images of proximal femur trabecular bone (left) and mid-diaphysis cortical bone (right). Graphs showing trabecular bone volume/tissue volume (BV/TV), trabecular number (Tb.N), trabecular thickness (Tb.Th), trabecular spacing (Tb.Sp), cortical thickness (Ct.Th), internal cortical diameter and cortical bone mineral density (BMD). (**c**) Graphs showing yield load, maximum load, fracture load, stiffness and energy dissipated prior to fracture derived from 3-point bend testing of femurs. (**d**) Graphs showing yield load, maximum load and stiffness derived from compression testing of vertebra. Scale bars: a, 1mm and b, 0.5mm.

**Figure S17**. Increased bone mineral content and strength in adult *DNA Regulator of Cell Cycle* knockout mice (*Rgcc^−/−^*) (a) X-ray microradiography images of femur and caudal vertebrae from female wild-type (WT) and *Rgcc^−/−^* mice at postnatal day 112 (P112). Graphs show reference ranges derived from 320 WT mice, mean (solid line), 1.0SD (dotted lines) and 2.0SD (grey box). Parameters from individual *Rgcc^−/−^* mice are shown as red dots and mean values as a black line (n=2 animals). (**b**) Micro-CT images of proximal femur trabecular bone (left) and mid-diaphysis cortical bone (right). Graphs showing trabecular bone volume/tissue volume (BV/TV), trabecular number (Tb.N), trabecular thickness (Tb.Th), trabecular spacing (Tb.Sp), cortical thickness (Ct.Th), internal cortical diameter and cortical bone mineral density (BMD). (**c**) Graphs showing yield load, maximum load, fracture load, stiffness and energy dissipated prior to fracture derived from 3-point bend testing of femurs. (**d**) Graphs showing yield load, maximum load and stiffness derived from compression testing of vertebra. Scale bars: a, 1mm and b, 0.5mm.

**Figure S18**. Increased bone mass and brittle bones in adult *Tyrosine 3-Monooxygenase/Tryptophan 5-Monooxygenase Activation Protein Epsilon* knockout mice (*Ywhae^−/−^*) (**a**) X-ray microradiography images of femur and caudal vertebrae from female wild-type (WT) and *Ywhae^−/−^* mice at postnatal day 112 (P112). Graphs show reference ranges derived from 320 WT mice, mean (solid line), 1.0SD (dotted lines) and 2.0SD (grey box). Parameters from individual *Ywhae^−/−^* mice are shown as red dots and mean values as a black line (n=2 animals). (**b**) Micro-CT images of proximal femur trabecular bone (left) and mid-diaphysis cortical bone (right). Graphs showing trabecular bone volume/tissue volume (BV/TV), trabecular number (Tb.N), trabecular thickness (Tb.Th), trabecular spacing (Tb.Sp), cortical thickness (Ct.Th), internal cortical diameter and cortical bone mineral density (BMD). (**c**) Graphs showing yield load, maximum load, fracture load, stiffness and energy dissipated prior to fracture derived from 3-point bend testing of femurs. (**d**) Graphs showing yield load, maximum load and stiffness derived from compression testing of vertebra. Scale bars: a, 1mm and b, 0.5mm.

**Figure S19**. Bone composition and structure analysis from rapid throughput screening murine knockouts. (**a**) The graphs demonstrate the relationship between bone mineral content and yield load, maximum load, fracture load and stiffness in femurs from P112 female WT mice (N = 320). For yield load, maximum load, and stiffness the blue line shows the linear regression (P = 0.005, P < 0.00001, P = 0.003 respectively) and the grey box indicates ± 2SD. For fracture load the blue line shows the linear regression (P = 0.00003) and the grey box indicates ± 95% confidence intervals. The mean values for *Cbx1^+/−^, Dscc1^+/−^, Rgcc^−/−^, Wac^+/−^* and *Ywhae^−/−^* (N = 2 from OBCD screen) mice are shown in orange. The *Wac^+/−^* femur yield load was 2.8 SD above the wild type reference range and *Dscc1^+/−^* fracture load was on the 1.7^th^ centile. (**b**) The graph demonstrates the relationship between bone mineral content and yield load, maximum load and stiffness in vertebrae from P112 female WT mice (N = 320). For yield and maximum loads the blue line shows the linear regression (P = <0.00001) and the grey box indicates ± 95% confidence intervals. For stiffness the blue line shows the linear regression (P = 0.0001) and the grey box indicates ± 2SD. The mean values for *Cbx1^+/−^, Dscc1^+/−^, Rgcc^−/−^, Wac^+/−^* and *Ywhae^−/−^* (N = 2 from OBCD screen) mice are shown in orange.

**Figure S20. Bivariate scatterplots describing pairwise comparisons of each of the first 20 ancestry informative principal components derived from unrelated subjects of the 1000 Genomes study**. Data points represent subjects that are coloured according to their predefined 1000 genomes study population*. Pairwise combinations involving eigenvectors 1,2 and 5 represented the smallest number of eigenvectors that were able to adequately resolve the British sub-population (GBR) from other ethnicities and were subsequently used to for clustering and ancestry assignment of the UK Biobank sample. *CHB=Han Chinese in Beijing, China, JPT=Japanese in Tokyo, Japan, CHS=Southern Han Chinese, CDX=Chinese Dai in Xishuangbanna, China, KHV=Kinh in Ho Chi Minh City, Vietnam, CEU=Utah Residents (CEPH) with Northern and Western European Ancestry,TSI=Toscani in Italia, FIN=Finnish in Finland, GBR=British in England and Scotland, IBS=Iberian Population in Spain, YRI=Yoruba in Ibadan, Nigeria, LWK=Luhya in Webuye, Kenya, GWD=Gambian in Western Divisions in the Gambia, MSL=Mende in Sierra Leone, ESN=Esan in Nigeria, ASW=Americans of African Ancestry in SW USA, ACB=African Caribbeans in Barbados, MXL=Mexican Ancestry from Los Angeles USA, PUR=Puerto Ricans from Puerto Rico, CLM=Colombians from Medellin, Colombia, PEL=Peruvians from Lima, Peru, GIH=Gujarati Indian from Houston, Texas, PJL=Punjabi from Lahore, Pakistan, BEB=Bengali from Bangladesh, STU=Sri Lankan Tamil from the UK, ITU=Indian Telugu from the UK

**Figure S21. Evaluating expectation maximization clustering model fit**. The number of predefined clusters is described on the x-axis and model fit on the y-axis [Inferred by three model selection criteria: i.e. log-likelihood (LogL), Akaike information criteria (AIC), and Bayesian information criterion (BIC)]. Twelve predefined clusters were chosen for clustering as sensitivity analyses suggested that this number provided a good compromise between model fit and computational burden (i.e. more clusters requires more computation).

**Figure S22. Bivariate scatterplots describing pairwise comparisons of ancestry informative principal components 1,2 and 5 derived from unrelated subjects of the 1000 genomes study and all subjects from the UK Biobank sample**. Data points represent subjects that are coloured according to their allocated cluster, as estimated by Expectation Maximization (EM) clustering. Samples from the UK-Biobank sample are annotated using “UKB”. Other 1000 genomes poulations are annotated using the following: CHB=Han Chinese in Beijing, China, JPT=Japanese in Tokyo, Japan, CHS=Southern Han Chinese, CDX=Chinese Dai in Xishuangbanna, China, KHV=Kinh in Ho Chi Minh City, Vietnam, CEU=Utah Residents (CEPH) with Northern and Western European Ancestry,TSI=Toscani in Italia, FIN=Finnish in Finland, GBR=British in England and Scotland, IBS=Iberian Population in Spain, YRI=Yoruba in Ibadan, Nigeria, LWK=Luhya in Webuye, Kenya, GWD=Gambian in Western Divisions in the Gambia, MSL=Mende in Sierra Leone, ESN=Esan in Nigeria, ASW=Americans of African Ancestry in SW USA, ACB=African Caribbeans in Barbados, MXL=Mexican Ancestry from Los Angeles USA, PUR=Puerto Ricans from Puerto Rico, CLM=Colombians from Medellin, Colombia, PEL=Peruvians from Lima, Peru, GIH=Gujarati Indian from Houston, Texas, PJL=Punjabi from Lahore, Pakistan, BEB=Bengali from Bangladesh, STU=Sri Lankan Tamil from the UK, ITU=Indian Telugu from the UK.

**Figure S23. Targeting DAAM2 exon 2 with CRISPR/Cas9 induced double stranded breaks reduced DAAM2 protein level in SaOS-2 cells. A**) DAAM2 protein level quantification in control cells and edited DAAM2 cells (gRNA1 and gRNA2). Bars represent the mean of six independent experiments ± SEM. *** represent P < 0.001 compared to control cells determined by one-way Anova and Bonferroni post-hoc tests. **B**) Bands from representative Western Blots of DAAM2 (upper panel) and total protein (lower panel) of at least six independent experiments from different cell line passages. Ct: controls; gRNA1: *DAAM2* edited cells with gRNA1; gRNA2: *DAAM2* edited cells with gRNA2.

## Acknowledgments

This research has been conducted using the UK Biobank Resource (accession IDs: 24268, 12703 and 4580). J.B.R. was supported by the Canadian Institutes of Health Research, the Canadian Foundation for Innovation and the Fonds de Recherche Santé Québec (FRSQ) and a FRQS Clinical Research Scholarship. TwinsUK is funded by the Wellcome Trust, Medical Research Council, European Union, the National Institute for Health Research (NIHR)-funded BioResource, Clinical Research Facility and Biomedical Research Centre based at Guy’s and St Thomas’ NHS Foundation Trust in partnership with King’s College London. J.A.M was funded by the Canadian Institutes of Health Research. D.M.E. was funded by a National Health and Medical Research Council Senior Research Fellowship (APP1137714) and funded by a Medical Research Council Programme Grant (MC_UU_12013/4). J.P.K was funded by a University of Queensland Development Fellowship. CLG was funded by Arthritis Research UK (ref; 20000). G.R.W., J.H.D.B. and P.I.C. were funded by the Wellcome Trust (Strategic Award grant number 101123; project grant 094134) and P.I.C was also funded by the Mrs Janice Gibson and the Ernest Heine Family Foundation. D.K. was supported by Israel Science Foundation grant #1283/14. Y.-H.H. was funded by US NIH NIAMS 1R01AR072199. F.R., C.M.-G., and K.T. were funded by the Netherlands Organization for Health Research and Development (ZonMw VIDI 016.136.361 grant). C.L.A.-B. was funded by NIH/NIAMS AR063702 AR060981. D.P.K. was funded by grants from the National Institute of Arthritis Musculoskeletal and Skin Diseases R01 AR041398, R01 AR072199. S.Y. was funded by the Australian Government Research Training Program Scholarship. J.R. and S.K. were funded by the Genetic Factors of Osteoporosis-GEFOS EU FP7 Integrated Project Grant Reference: 201865 2008-12 and 2007-12 UK NIHR Biomedical Research Centre Grant (Musculoskeletal theme) to Cambridge Clinical School. C.O. was supported by the Swedish Research Council, Swedish Foundation for Strategic Research, ALF/LUA research grant from the Sahlgrenska University Hospital, Lundberg Foundation, European Calcified Tissue Society, Torsten and Ragnar Söderberg’s Foundation, Novo Nordisk Foundation, Knut and Alice Wallenberg Foundation.

We thank M. Schull for assistance with high-performance computing at the University of Queensland Diamantina Institute, and T. Winkler for invaluable technical support for the EasyStrata Software used in this study. We thank the Sanger Institute’s Research Support Facility, Mouse Pipelines and Mouse Informatics Group who generated the mice and collected materials for this manuscript. We would like to thank the research participants and employees of 23andMe for making this work possible. The following members of the 23andMe Research Team contributed to this study: Michelle Agee, Babak Alipanahi, Adam Auton, Robert K. Bell, Katarzyna Bryc, Sarah L. Elson, Pierre Fontanillas, Nicholas A. Furlotte, Jennifer C. McCreight, Karen E. Huber, Nadia K. Litterman, Matthew H. McIntyre, Joanna L. Mountain, Elizabeth S. Noblin, Carrie A.M. Northover, Steven J. Pitts, J. Fah Sathirapongsasuti, Olga V. Sazonova, Janie F. Shelton, Suyash Shringarpure, Chao Tian, Joyce Y. Tung, Vladimir Vacic, and Catherine H. Wilson.

